# Glycoengineering of NK cells with glycan ligands of CD22 and selectins for B-cell lymphoma therapy

**DOI:** 10.1101/2020.03.23.004325

**Authors:** Senlian Hong, Chenhua Yu, Peng Wang, Yujie Shi, Bo Cheng, Mingkuan Chen, Digantkumar G. Chapla, Natalie Reigh, Yoshiki Narimatsu, Xing Chen, Henrik Clausen, Kelly W. Moremen, Matthew Scott Macauley, James C. Paulson, Peng Wu

## Abstract

CD22, a member of Siglec family of sialic acid binding proteins, has restricted expression on B cells. Antibody-based agents targeting CD22 or CD20 (Rituxan™) on B lymphoma and leukemia cells exhibit clinical efficacy for treating these malignancies, but also attack normal B cells leading to immune deficiency. Here, we report a chemoenzymatic glycocalyx editing strategy to introduce high-affinity and specific CD22 ligands onto NK-92MI and cytokine-induced killer (CIK) cells to achieve tumor-specific CD22 targeting. These CD22-ligand modified cells exhibited significantly enhanced tumor cell binding and killing *in vitro* without harming healthy B cells. For effective lymphoma cell killing in vivo we further functionalized CD22 ligand-modified NK-92MI cells with the E-selectin ligand sialyl Lewis X to promote trafficking to bone marrow. The cells containing the ligands of both CD22 and selectins resulted in the efficient suppression of B lymphoma in a xenograft model. Our results suggest that NK cells modified with glycan ligands to CD22 and selectins promote both targeted killing of B lymphoma cells and improved trafficking to sites where the cancer cells reside, respectively.

## Introduction

Current treatments for patients with B-cell malignancies, such as non-Hodgkin lymphoma and leukemia primarily rely on systemic chemotherapy and the anti-CD20 antibody rituximab. However, when people become resistant to both, new treatments are needed.^1,2^ CD22, also known as sialic acid-binding immunoglobulin-like lectin 2 (Siglec-2), is a B-cell restricted inhibitory receptor for antagonizing B cell receptor (BCR) signaling. CD22 is highly expressed by various B lymphoma and leukemia.^3,4^ Its B-cell restricted expression and endocytic nature makes CD22 a promising therapeutic target. CD22-targeting antibodies and antibody-drug conjugates based on Anti-CD22 based therapies have shown clinical efficacies to treat aggressive and follicular non-Hodgkin’s lymphoma, acute lymphoblastic leukemia and diffuse large B cell lymphoma.^5^ However, these agents cannot differentiate healthy B cells from B malignancies and cause serious side effects such as low blood counts, systemic cytotoxic effects and veno-occlusive disease (VOD). Recently, Fry and co-workers reported that anti-CD22 engineered Chimeric antigen receptor T cells (CAR-Ts) induced similar response rates as anti-CD19-CAR-T cells in patients with acute lymphoblastic leukemia.^6–9^ However, CAR-T therapies face the same dilemma of on-target-off-tumor toxicity.

Unnatural sialic acid analogs with high-affinity and specificity for CD22 are promising surrogates of antibodies for CD22 targeting.^10–14^ By introducing unnatural substituents onto C-9 of Neu5Ac, sialic acid analogs with micro-molar affinity for CD22 have been discovered, such as 9-*N*-biphenylcarboxamide-Neu5Ac (^BPC^Neu5Ac^10^; **1**) and 9-*N*-*m*-phenoxybenzamide-Neu5Ac (^MPB^Neu5Ac^12^; **2**). Nanoparticles, fabricated using synthetic glycolipids, that display these ligands multivalently have been used to deliver cytotoxic drugs to CD22 over-expressed tumor cells in animal models.^15–17^Like antibody based reagents, nanoparticles equipped with CD22 ligands target all cells expressing CD22. To employ CD-22 targeting ligands to channel cytotoxic agents for tumor-specific killing, the challenge is to de-couple the killing mechanism from the CD-22 targeting.

NK-92MI is a non-immunogenic and constantly cytotoxic NK cell line currently undergoing intensive clinical trials as an “off-the-shelf therapeutic” for treating both hematological and solid malignancies.^18–20^ NK-92MI-induced killing relies on the interaction of NK activation receptors, e.g. NKG2D, with their ligands that have restricted expressions on stressed and malignant cells, and thus systemic toxicity is eliminated.^20^ Cytokine-Induced Killer (CIK) cells, generated by ex vivo incubation of human peripheral blood mononuclear cells (PBMC) or cord blood mononuclear cells, feature a mixed T-and natural killer (NK) cell-like phenotype.^21–24^ Like NK-92MI cells, the intrinsic tumor killing ability of CIK is mainly mediated by the NKG2D receptor that recognizes, in MHC-independent manner, stress-inducible targets.^24^ We hypothesize that by using high-affinity and specific CD22 ligands to functionalize NK-92MI and CIK cells, specific targeting and killing of tumor cells with CD22 over-expression can be achieved.^25,26^

Here, we exploit chemoenzymatic glycan editing^27^ to create high-affinity CD22 ligands on the cell surface of NK-92MI and CIK cells. We demonstrate that the functionalized NK-92MI and CIK cells display high-affinity CD22 ligands on their N-glycans and form multivalent ligand presentation, which leads to significantly enhanced binding and killing of CD22 over-expressed tumor cells *in vitro* while leaving healthy B cells untouched. *In vivo* in a xenograft lymphoma model, however, the functionalized NK-92MI cells only show weak activities. In this model, E-selectin that is constitutively expressed in the bone marrow binds to lymphoma cells, facilitates their bone-marrow deposition, and shed them from attacking by NK-92MI cells. We discovered that NK-92MI cells that are functionalized with CD22 ligands can be further engineered via fucosyltransferase-mediated installation of E-selectin ligands to enhance their bone marrow migration. In this way, significant suppression of B-lymphoma proliferation *in vivo* is achieved.

## Results

### Engineering CD22 ligands on live cells via chemoenzymatic glycan editing

To engineer killer cells for targeting CD22 positive tumors, we sought to install CD22 ligands onto their cell surfaces directly. Because the sialylated epitope Neu5Acα2-6Galβ1-4Glc*N*Ac found at peripheral *N*-glycans is a natural ligand of CD22,^14^ we first evaluated the possibility of employing sialyltransferase (ST)-mediated sialylation to create this epitope on mammalian cell surfaces by using Chinese hamster ovary (CHO) Lec2 mutant cells^28,29^ and an engineered HEK293 cell line^30^ (HEK293-∆ST), a human epithelial cell line with inactivated ST6Gal1/2 and ST3Gal3/4/6 sialyltransferase genes and selective loss of sialic acids on N-glycans and glycolipids but not some types of O-glycans, as model systems. Both cell lines possess complex type *N*-glycans terminated with *N*-acetyllactosamine (type 2 Lac*N*Ac Galβ1-4Glc*N*Ac, or in the case of HEK293 also minor amounts of LacDi*N*Ac, Gal*N*Acβ1-4Glc*N*Ac) as is the case for most mammalian cells.

Two α-2,6STs, the truncated human 2,6ST (ST6Gal1^31^) and *Photobacterium damsela* 2,6ST (Pd2,6ST^32,33^), were evaluated for this endeavor using biotinylated CMP-Neu5Ac^34^ as the donor. Approximately 2-fold more biotin-Neu5Ac was incorporated onto Lec2 cells by ST6Gal1 compared to that added by Pd2,6ST (Supplementary figure S1). Therefore, ST6Gal1 was chosen for the follow-up cell-surface glycan engineering experiment (Figure 1a).

**Figure 1.**
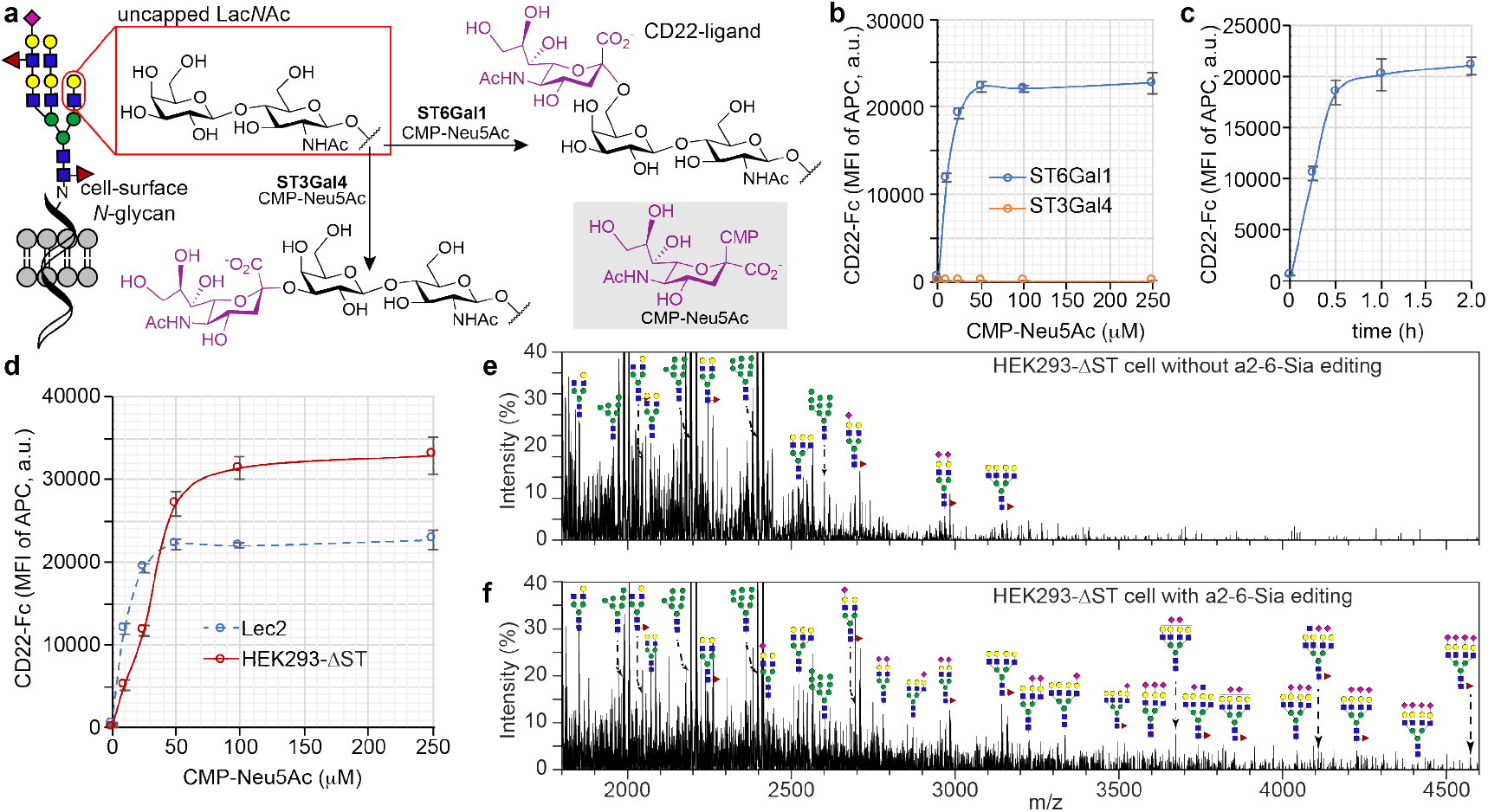
Chemo-enzymatic generation of sialyl ligands on live cells to increase CD22 binding. (a) ST6Gal1- and ST3Gal4-assisted incorporation of Neu5Ac onto the uncapped type 2 Lac*N*Ac in α2-6- and α2-3-linkages, respectively. Among the resulting epitopes, Neu5Acα2-6Galβ1-4Glc*N*Ac showed high affinity against CD22, while the Neu5Acα2-3Galβ1-4Glc*N*Ac did not. (b-d) Flow cytometry quantification of CD22-Fc binding onto ST-engineered live cells. The error bars represent the standard deviation of three biological replicates. (b) CD22-Fc binding of ST6Gal1 or ST3Gal-modified Lec2 cells under different doses of CMP-Neu5Ac substrate. (c) CD22-Fc binding of ST6Gal1-modified Lec2 cells for different periods. (d) Comparison of CD22-Fc binding of Lec2 and HEK293-∆ST mutant cells modified by ST6Gal1. (e and f) The matrix-assisted laser desorption/ionization time-of-flight (MALDI-TOF) mass spectrometer based resolution of the profile of *N*-glycans in HEK293T-∆ST mutant cells without (e) or with 2,6ST-engineering (f).

Incubating Lec2 and HEK293-∆ST cells with ST6Gal1 in the presence of CMP-Neu5Ac resulted in robust CD22-Fc binding, whereas the cells treated with ST3Gal4^35,36^ and CMP-Neu5Ac to create Neu5Acα2-3Galβ1-4Glc*N*Ac on the surface only exhibited background signals (Figure 1b and c and Supplementary figure S3a). As revealed by matrix-assisted laser desorption/ionization time-of-flight mass spectrometer (MALDI-TOF) (Figure 1e), after ST6Gal1 mediated sialylation, sialylated di-, tri-, and tetra-branched *N*-glycans could be detected on the modified HEK293-∆ST cells (Figure 1f). Interestingly, under saturated conditions a comparable level of newly added-Neu5Ac on Lec2 and HEK293-∆ST cells (Supplementary figure S4) led to a 1.5-fold higher CD22-Fc binding on HEK293-∆ST cells than on Lec2 cells (Figure 1d).

Two unnatural Neu5Ac analogs, ^BPC^Neu5Ac (**2a**) and ^MPB^Neu5Ac (**2b**), when α-2,6-linked to LacNAc, are known to bind to CD22 with affinities in the submicromolar range.^12^ We then assessed the feasibility of using the corresponding CMP sugars of these two analogs as the donor substrates for ST6Gal1-assisted sialylation to generate high-affinity CD22 ligands on the cell surface (Figure 2a, Supplementary figure S16, and s17). At all donor concentrations assessed, Lec2 cells (Figure 2b and Supplementary figure S5a) and HEK293-∆ST cells (Supplementary figure S5b-d) that were modified with unnatural substrates showed significantly higher binding to CD22-Fc binding than the cells treated with CMP-Neu5Ac. Under saturating conditions, approximately 10-fold and 500-fold higher CD22 binding was detected on CMP-^BPC^Neu5Ac- and CMP-^MPB^Neu5Ac-modified cells than the cells treated with and without natural CMP-Neu5Ac, respectively. ^BPC^Neu5Ac and ^MPB^Neu5Ac installed on the cell surface in the α2-3-linkage only exhibited very weak CD22-Fc binding on Lec2 cells (Figure 2c and Supplementary figure S5c) and HEK293-∆ST cells (Supplementary figure S5e and s5f). Notably, under saturating conditions approximately 2-fold higher CD22-Fc binding was detected on **2a** and **2b** modified HEK293-∆ST cells than their Lec2 counterparts, respectively, (Figure 2d). This observation suggests that cell-specific glycan scaffolds may influence binding to CD22 ligands engineered on the surface.

**Figure 2.**
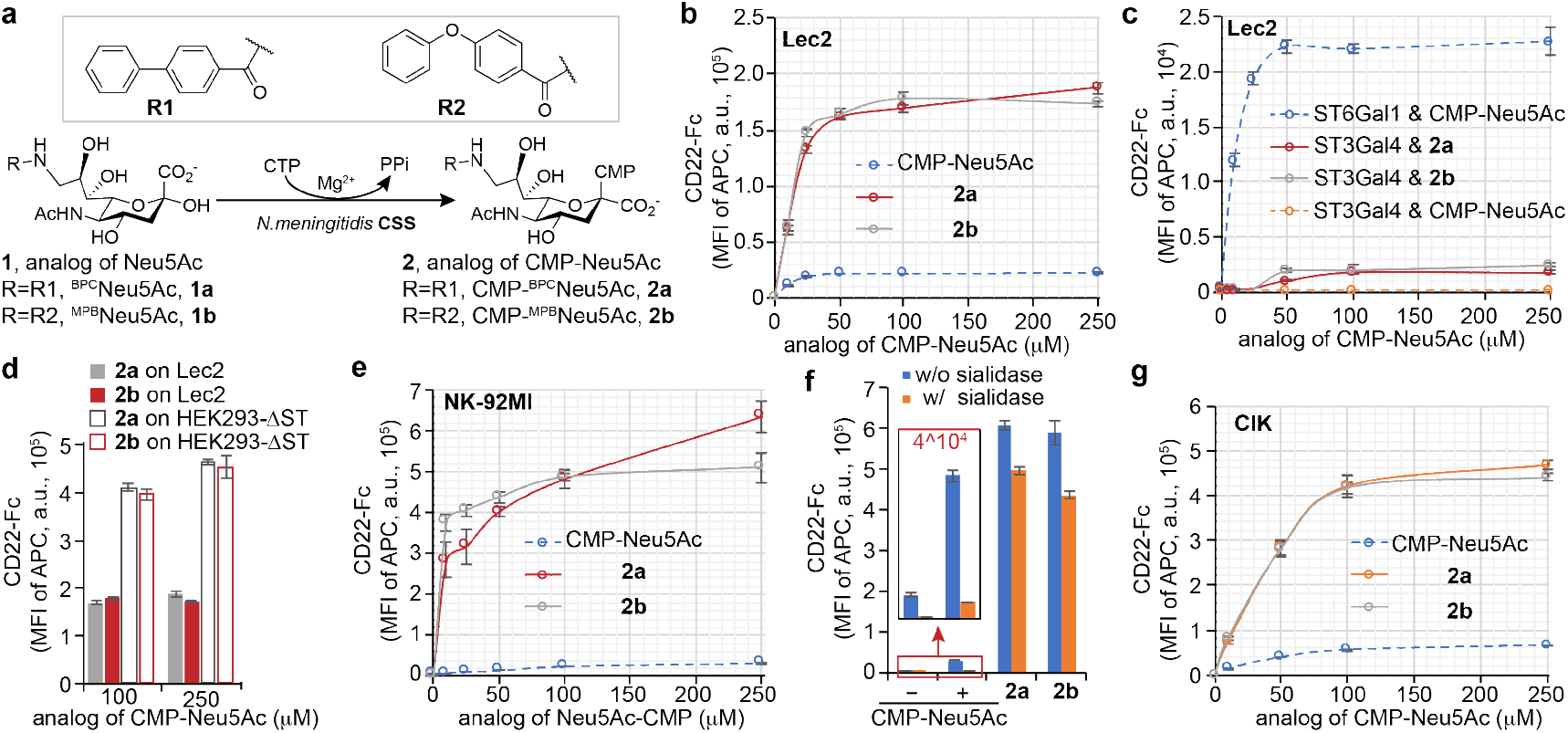
hSTs-assisted creation of high-affinity CD22-ligands on live cells. (a) synthesis of C-9 functionalized analogs of CMP-Neu5Ac, including CMP-^BPC^Neu5Ac (**2a**) and CMP-^BPC^Neu5Ac (**2b**). (b-g) flow cytometry quantification of CD22-Fc binding onto STs-engineered live cells, including Lec2, HEK293-∆ST, NK-92MI and CIK cells. The error bars represent the standard deviation of three biological replicates. The CD22-Fc based probing of the ST6Gal1-(b) or ST3Gal4-assisted (c) incorporation of unnatural sialic acids (**2a** or **2b**) or nature sialic acid (Neu5Ac) onto Lec2 cells. (d) Comparison of CD22-Fc binding of Lec2 and HEK293-∆ST mutant cells modified by ST6Gal1 and unnatural sialic acids. (e and f) The CD22-Fc based probing of ST6Gal1-assisted incorporation of the different concentrations of **2a**, **2b**, or Neu5Ac onto NK-92MI cells (e) or NK-92MI cells pretreated by sialidase or not (f). (g) The CD22-Fc based probing of the ST6Gal1-assisted incorporation of sialic acids onto CIK cells.

### Engineering high-affinity ligands on NK-92MI and CIK cells for efficient CD22-targeting

After confirming that **2a** and **2b** can be incorporated onto mammalian cell surfaces via ST6Gal1-mediated sialylation to create high-affinity CD22 ligands, we then explored the feasibility of using this strategy to engineer NK-92MI and CIK cells for targeting tumor cells expressing CD22. To prepare CIK cells, peripheral blood precursors from healthy donors were activated with anti-CD3 and anti-CD28 antibodies, and maintained in IL-2 supplemented T-cell culture medium for 10–15 days.^24^ After subjecting NK-92MI and CIK cells to ST6Gal1-mediated sialylation, both cell lines exhibited dramatically enhanced binding to CD22-Fc in a dose- and time-dependent manner (Figure 2e, 2g, Supplementary figure S6a, and s7a). Under saturating concentrations of CMP-^BPC^Neu5Ac and CMP-^MPB^Neu5Ac, the high affinity ligand modified NK-92MI cells showed approximately 20–50-fold higher CD22 binding in comparisons to their counterparts treated with CMP-Neu5Ac to install the natural ligands. Likewise, CMP-^BPC^Neu5Ac- and CMP-^MPB^Neu5Ac-modified CIK cells showed approximately 10-fold higher CD22 binding compared to their counterparts treated with CMP-Neu5Ac. We confirmed that the newly created CD22 ligands did not impair the proliferation and cell viability of NK-92MI and CIK cells (Supplementary figure S6c-e, and s7c). Three days after *in vitro* sialylation, even with proliferation- or turnover-associated dilution of cell-surface ligands the modified NK-92MI and CIK cells still showed approximately 10- and 15-fold better CD22-Fc binding than the untreated cells, respectively (Supplementary figure S6f and s7b). Desialylation by neuraminidase treatment followed by enzymatic sialylation of NK-92MI and CIK cells did not increase CD22 binding (Figure 2f, Supplementary figure S6b, s7h and s7i), but rather impaired CD22 binding especially when the natural donor substrate CMP-Neu5Ac was used for sialylation. This observation suggests that in the natural scenario, Neu5Acα2-6Galβ1-4Glc*N*Ac on O-glycans and glycolipid also contribute significantly toward CD22 binding. However, when the high-affinity CD22 ligands are incorporated onto N-glycans, the contributions of the natural ligands on O-glycans and glycolipid become less important.

### Killer cells armed with CD22-ligands exhibit significantly enhanced binding and killing of B-lymphoma cells in vitro

Enhanced CD22-Fc binding of the ligand functionalized NK-92MI or CIK cells should translate into their enhanced ability to induce the binding and killing of CD-22 over-expression tumor cells. The targeting efficiency of CD22-ligand-loaded NK-92MI and CIK for B-lymphoma cells with high CD22 expression was assessed by quantifying the formation of cell-cell conjugates when coculturing NK-92MI or CIK (effector cells, E) and B-lymphoma cells (target cells, T). In comparison to the unmodified counterparts, the CD22-ligand modified NK-92MI and CIK significantly increased the formation of cell-cell conjugates, respectively. The higher the level of CD22 expression of target cells (Figure 3a), the higher percentage of conjugate formation was detected (Figure 3b, 3c, Supplementary figure S7d, s7e, and s8). Modified Killer cells did not induce enhanced cluster formation with Jurkat cells that are CD22 negative. Consistent with the enhanced binding, the functionalized NK-92MI and CIK cells induced better killing of target B-lymphoma cells (Figure 3d-h, Supplementary figure S7f, and s7g), which was further supported by the increased IFN-gamma production (Supplementary figure S9).

Next, we tested CD22-targeting efficiency in the presence of natural ligands on other cells, like red blood cells (RBCs). RBC present nature ligands against CD22 on their surface, as shown by CD22-Fc binding results (Supplementary figure S10). Interestingly, CD22-ligand-mediated targeting of NK-92MI or CIK cells to B-lymphoma cells was 2-fold improved after docking 100-times of RBC cells into the coculture systems (Figure 3i, 3j and Supplementary figure S7d). Unlike killer cells, human RBCs do not have intrinsic cytotoxicity and cannot kill B-lymphoma cells. The presence of CD22 high-affinity ligand armed RBC did not kill Raji and Daudi cells (Supplementary figure S11). Moreover, while the modified NK-92MI cells bound to normal B cells from healthy donors, there was no significantly specific lysis, likely because the level of CD22 on normal B cells is only about 20% that found on Raji cells (Figure 3a and Supplementary figure S12). These results suggest that killer cells armed with CD22-ligands may be a biocompatible tool for selective eradication of B-lymphoma.

**Figure 3.**
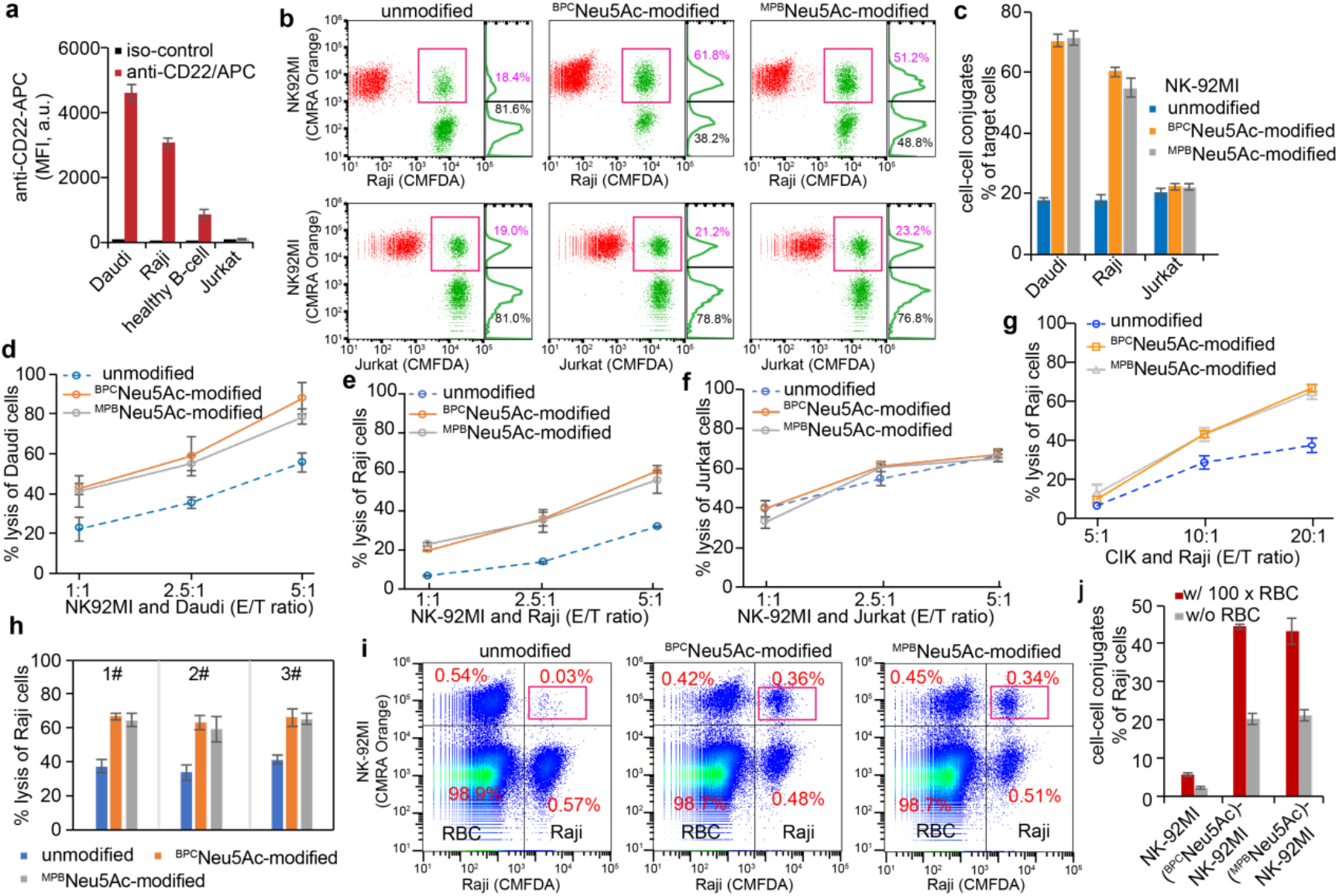
Targeting B-lymphoma with the CD22-sialoside-ligands armed killer cells. (a) Comparison of CD22 expression levels on different types of cells, including two B-lymphoma cell lines Daudi and Raji, B-cell from six healthy donors, as well as one T-lymphoma cell line Jurkat. (b) Flow cytometry-based detection of cell-cell conjugates of NK-92MI cells (effector cell, stained with CMRA Orange) and target cells (Raji or Jurkat cells, stained with CMFDA Green) at an E/T ratio of 5:1. Number (in purple) indicates the ratio of target cells (i.e., Raji or Jurkat cells) in conjugation with NK-92MI cells. (c) The relative population of NK-92MI cell-cell conjugates with target cells. (d-h) LDH release assay based quantifying of killer cellcytotoxicity against Daudi (d), Raji (e, g and h), and Jurkat cells (f). In figure d-f, NK-92MI was the effector cell. In figure g and h, CIK was the effector cell. (i and j) Flow cytometry-based detection of cell-cell conjugates of NK-92MI cells and Raji cells (E/T ratio, 5:1), after docking of 100 × excess human red blood cells (RBC) or not.

### Exploiting glycan engineering for efficient NK-92MI-immunotherapy in a mouse model

The antitumor efficacies of NK-92MI armed with high-affinity CD22-ligands were next evaluated in vivo using a xenograft model. Raji cells expressing firefly luciferase (Raji-Luc) were intravenously (*i.v.*) infused into NSG mice on day 0, followed by three NK-92MI injections (*i.v.*) on day 2, day 6, and day 10. The growth of tumors was monitored by longitudinal, non-invasive bioluminescence imaging (Figure 4a and 4b). Compared to the control group that was received buffer injection only, no significant inhibition of Raji-Luc tumor growth was observed in the group treated with unmodified NK-92MI cells, whereas ^BPC^Neu5Ac-NK-92MI cells exhibited moderate capabilities to suppress tumor growth with ~25% reduction in bioluminescence signals relative to the control group treated with the buffer only (Figure 4b).

**Figure 4.**
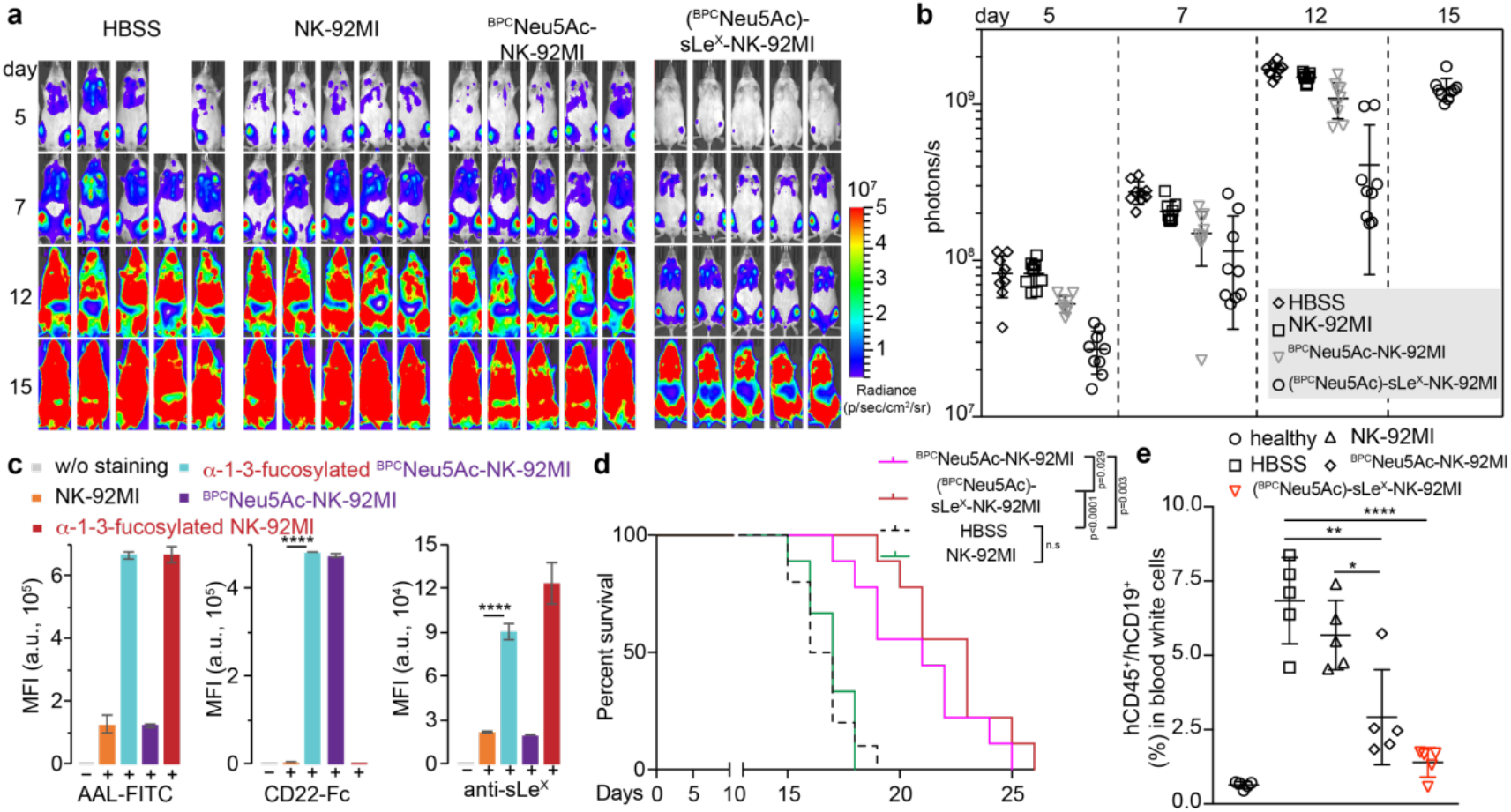
NK-92MI-based eradication of Raji B-lymphoma in a murine xenograft model. B-cell malignancies were established by tail vein injection of 1 million Raji-Luc cells into NSG mice on day 0. On day 2, the tumor-bearing mice received the 1^st^ treatment by i.v. infusion of 10 million NK-92MI cells. One group of mice received NK-92MI cells conjugated with ^BPC^Neu5Ac (^BPC^Neu5Ac-NK-92MI), and another group of mice received ^BPC^Neu5Ac-NK-92MI cells after the FuT6-assisted creation of sLe^X^ ((^BPC^Neu5Ac)-sLe^X^-NK-92MI); control groups received unmodified NK-92MI cells or a single systemic injection of the vehicle (HBSS). The 2^nd^ and 3^rd^ NK-92MI cell infusions were performed on day 6 and day 10, respectively. (a and b) On days 5, 7, 12, and 15, the mice were i.p. injected with D-luciferin and imaged by the IVIS system. (c) FuT6-assisted creation of sLe^X^ epitopes on NK-92MI cells. ^BPC^Neu5Ac was conjugated to NK-92MI in a α2-6-linkage by ST6Gal1, and cell-surface Neu5Acα2-3Galβ1-4GlcNAc was engineered into sLe^X^ after FuT6-assisted α1-3-fucosylation. The generation of CD22-ligands was assessed by CD22-Fc binding, the addition of α1-3-fucosylation was probed by AAL lectin staining, and the formation of sLe^X^ was confirmed by antibody staining. The error bars represent the standard deviation of three biological replicates. (d) Survival of mice after NK-92MI cell therapy illustrated by Kaplan-Meier curves. Shown are nine mice per treatment group pooled from three independent experiments. P values are calculated via Log-Rank (Mantel-Cox) test. (e) On day 16, the blood of tumor-bearing mice was collected, and the Raji cells (CD19^+^) in blood samples were analyzed with flow cytometry. * indicates the two-sided Students’ t-test p<0.05, **indicates the two-sided Students’ t-test p < 0.01, ****indicates the two-sided Students’ t-test p < 0.001.

Because most leukemia and lymphoma cells require a close relationship with the bone marrow microenvironment for their survival and disease progression.^37,38^ We reasoned that NK cells may not be sufficiently trafficking to bone marrow to attack B lymphoma where they reside *in situ*. The cell adhesion molecule E-selectin is constitutively expressed in the bone marrow and plays a critical role in promoting leukemia cell adhesion, survival, and resistance to chemotherapy. Consistent with this knowledge, we noticed that upon i.v. injection, the injected Raji cells primarily migrated to the bone marrow and the circulating Raji-Luc cells was only detectable in the bloodstream until late stages (14-16 days after Raji inoculation). By contrast, the majority of the injected NK-92MI cells were found in the lung, spleen and blood steam.^19,25^ Based on this observation, we hypothesized that increasing bone marrow infiltration would increase NK-induced tumor cell targeting and killing. The rolling and tethering of leukocytes on inflamed vessel endothelium cells rely on the binding of cell-surface sialyl Lewis X (sLe^X^, Neu5Acα2-3Galβ1-4(Fucα1-3)GlcNAc) with E-selectin to initialize tissue entry from the bloodstream, and importantly, cells migrate to bone through specialized marrow vessels that constitutively express vascular E-selectin. Therefore, we explored the possibility of creating sLe^X^ on the NK-92MI surface to enhance their interactions with E-selectin and thus facilitate bone migration.^39^ The similar strategy was employed previously by Sackstein and coworker to tune human multipotent mesenchymal stromal cell trafficking to bone.^40^

To create sLe^X^ on NK-92MI, the cells were treated with hFuT6^31,40,41^ and GDP-fucose. The successful fucosylation and creation of sLe^X^ on NK-92MI cells were confirmed by *Aleuria aurantia* lectin (AAL, specific for α1-3- and α1-6-linked fucose) and sLe^X^-specific antibody staining (Figure 4c). FT6-mediated sLe^X^ installation and ST6Gal1-mediated ^BPC^Neu5Asialylation could be performed simultaneously or sequentially without interfering each other. Moreover, NK-92MI cells displaying both sLe^X^ and CD22 ligands showed comparable Raji killing capabilities as ^BPC^Neu5Ac-NK-92MI cells (Supplementary figure S13). To assess if the newly created sLe^X^ promoted NK-92MI bone migration, dual-functionalized or ^BPC^Neu5Ac-modified NK-92MI cells were injected (*i.v.*) into NSG mice that had been inoculated with Raji-Luc for 12 day. The NK-92MI subsets in the periphery blood and the hind-leg bone marrow were quantified 10 hours later (Supplementary figure S14). Compared to ^BPC^Neu5Ac-NK-92MI cells without hFuT6-treatment, the bone marrow colonization of (^BPC^Neu5Ac)-sLe^X^-NK-92MI increased approximately 60% (Supplementary figure S14d) with a concomitant decrease of circling (^BPC^Neu5Ac)-sLe^X^-NK-92MI in the bloodstream (Supplementary figure S14c).

In contrast to the NK-92MI cells modified with CD22-ligand alone, these dual-functionalized NK-92MI cells showed remarkably enhanced capabilities to control Raji cell growth compared to ^BPC^Neu5Ac-NK-92MI (Figure 4a, 4b, 4d, and 4e). The lifespan of tumor-bearing NSG mice was extended approximately 27% in the (^BPC^Neu5Ac)-NK-92MI cell treated group and 40% in (^BPC^Neu5Ac)-sLe^X^-NK-92MI cell treated group, respectively, compared to the control group treated with buffer only.

## Conclusion and Discussion

Molecules presented on the cell surface determine how cells interact with their partners and their environment. By engineering the cell-surface landscape, cells can be endowed with the desired properties. In our previous studies^42^, by exploiting the donor substrate promiscuity of *H. pylori* fucosyltransferase, we demonstrated that HER2-specific antibody Herceptin were efficiently installed onto the cell-surface of NK-92MI cells. The resulting Herceptin-NK-92MI conjugates exhibited remarkably enhanced activities to induce the lysis of HER2-positve cancer cells both *ex vivo* and in a human tumor xenograft model. In the current study, we utilized ST6Gal1-mediated sialylation to create high-affinity CD22 ligands directly on NK-92MI cells. Via this approach significantly higher levels of CD22 ligands could be installed onto the cell surface in comparison to metabolic oligosaccharide engineering (MOE)-based ligand incorporation conducted by the Huang group^43^ (Supplementary figure S15). It is apparent that there is sufficiently high level of endogenous un-sialylated Lac*N*Ac residues that can be readily modified by ST6Gal1 to create a significant level of CD22 ligands. By contrast, the unnatural sialic acids (e.g., **1a** and **1b**) when added to the culture media for incorporation through the metabolic pathway can be inefficient as a result of competing with the natural sialic acid (Neu5Ac) for *de novo* sialylation of glycoproteins.

Major efforts on cancer immunotherapy are focused on the discovery of new tumor-specific antigens and ways to target them specifically. Relatively less attention has been devoted to approaches to facilitate the trafficking and tumor infiltration of immune cells. In their pioneering work,^40^ Sackstein, Xia et al., applied chemoenzymatic glycan editing based on human α1-3-fucosyltransferase (FT) to create sLe^X^ on human multipotent mesenchymal stromal cells and cord blood cells so as to enhance their engraftment and trafficking. Although a previous study introducing CD22 ligands into NK cells resulted in NK cell killing of B lymphoma cells as shown here,^43^ we found that equipping NK-92MI cells with selectin ligands in addition to CD22 ligands enhanced their efficacy in a B cell lymphoma model, where Raji B-lymphoma cells are localize in the bone marrow. For this reason, we suggest that the efficient trafficking of NK-92MI cells to bone marrow via engineered selectin ligands is critical for inducing productive tumor control.

In summary, cell-surface chemoenzymatic glycan editing offers an easy-to-practice strategy to boost the efficacy of therapeutic cells for better cancer treatment. Despite glycan turn-over and cell proliferation, a significant portion of CD22 ligands still remain on the cell surface after 72 hrs, which is in line with the persistence time of irradiated NK-92MI cells in human patients. Therefore, this technique provides a nice complement or synergistic approach to the permanent, genetic engineering approach that has found great success in constructing NK-92MI-based CAR-NK cells.

## Acknowledgements

Financial support was from the NIH (R01GM113046, R01GM111938, and R01GM093282 to P.W., P41GM103390, P01GM107012, and R01GM130915 to K.W.M., R01AI050143, P01HL107151, and U19Ai136443 to J.C.P.), and from the Danish National Research Foundation (DNRF107 to H.C. and Y.N.)

## Supporting Information

### Supporting Figures

**Figure S1.**
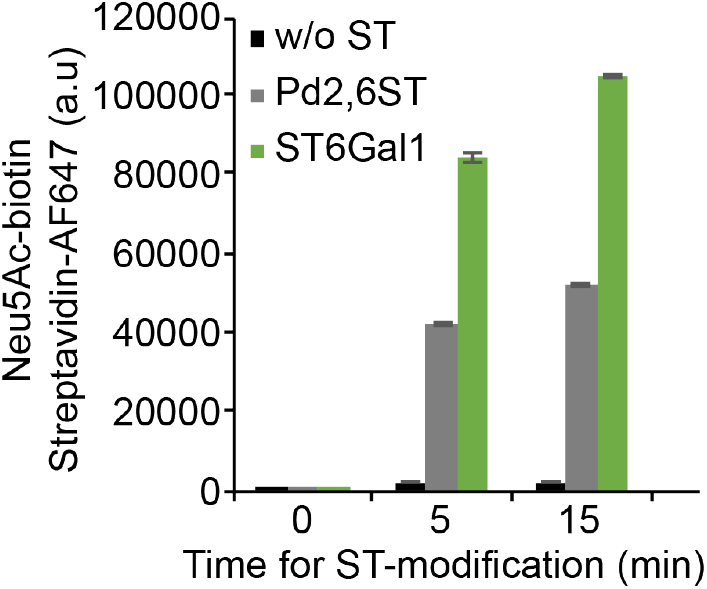
Chemoenzymatic incorporation of CMP-Neu5Ac-biotin onto Lec2 cell-surface glycans at α2-6-linkages. Lec2 cells were incubated in HBSS buffer (pH 7.4) containing 40 μg/mL STs (Pd2,6ST, ST6Gal1, or without ST), 100 μM CMP-Sia*N*Az-biotin, 3 mM HEPES, and 20 mM MgSO_4_ for the depicted time at 37 °C. After three times washing with 1 × DPBS (pH 7.4, precolded at 4 °C), the resulting cell-surface biotin was further probed with 2 μg/mL Streptavidin Alexa Fluor 647 (AF647) and quantified via flow cytometry. Error bars represent the standard deviation of three biological replicates.

**Figure S2.**
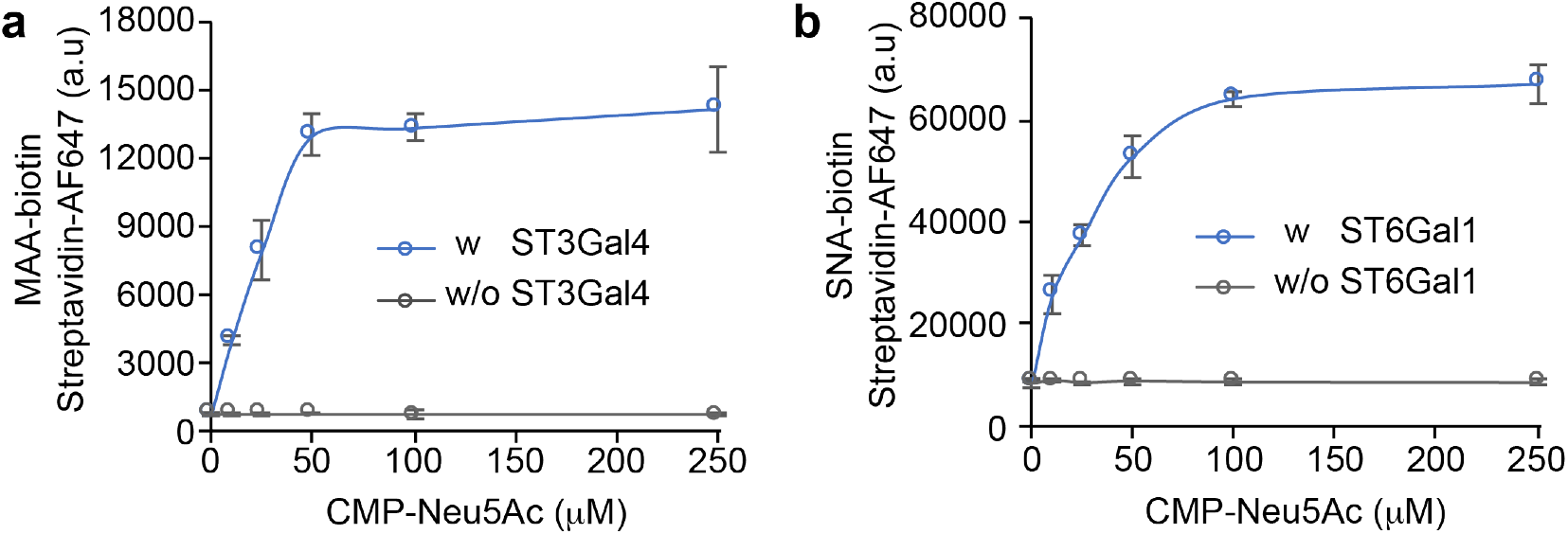
Lectin profiling of STs-assisted incorporation of sialic acids on live Lec2 cells at α2-3- or α2-6-linkages, respecitively. Lec2 cells were incubated in HBSS buffer (pH 7.4) containing 3 mM HEPES, 20 mM MgSO_4_, depicted concentration of CMP-Neu5Ac, and 40 μg/mL ST3Gal4 (a) or ST6Gal1 (b) for 30 min at 37 °C. After three times washing with 1 × DPBS (pH 7.4, precolded at 4 °C), the cell-surface sialic acids were detected with 20 μg/mL lectins (MAA-biotin for α2-3-linked sialic acids and SNA-biotin for α2-6-linked sialic acids). The resulting biotin was then labeled with 2 μg/mL Streptavidin-AF647 and quantified via flow cytometry. Error bars represent the standard deviation of three biological replicates.

**Figure S3.**
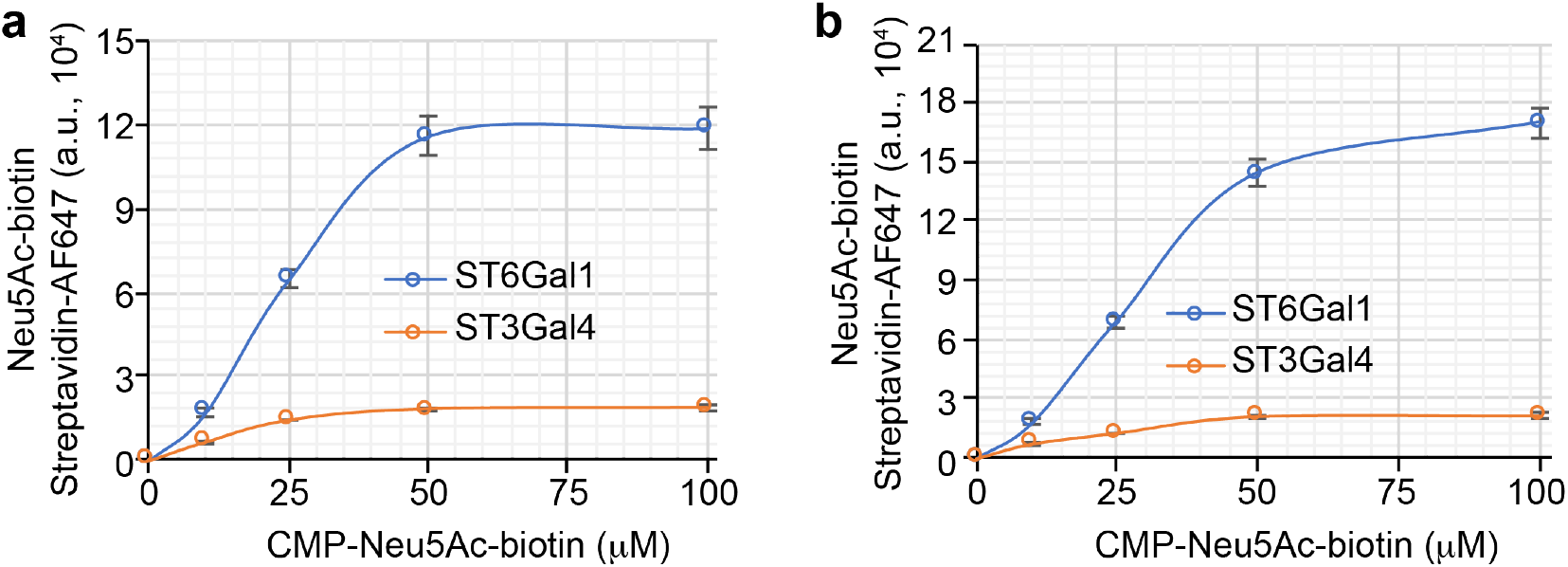
ST6Gal and ST3Gal4-assisted chemoenzymatic incorporation of CMP-Neu5Ac-biotin onto cell surface Nglycans. (a) Lec2 and (b) HEK293T-∆ST cells were incubated in HBSS buffer (pH 7.4) containing 40 μg/mL STs (ST6Gal1 or ST3Gal4), 3 mM HEPES, 20 mM MgSO_4_, and the depicted concentrations of CMP-Neu5Ac-biotin, for 30 min at 37 °C. After three times washing with 1 × DPBS (pH 7.4, precolded at 4 °C), the resulting cell-surface biotin was further probed with 2 μg/mL Streptavidin Alexa Fluor 647 (AF647) and quantified via flow cytometry. Error bars represent the standard deviation of three biological replicates.

**Figure S4.**
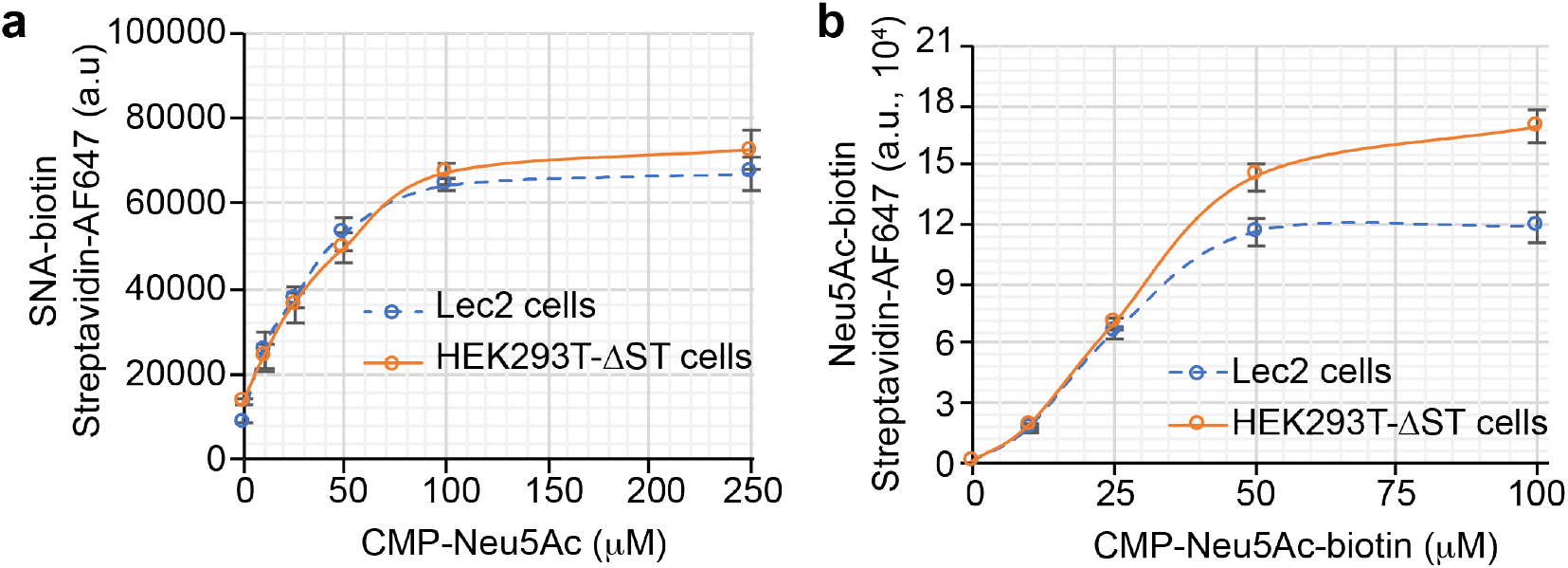
Profiling the α2-6-linked sialic acids incorporated by ST6Gal1 on live cells. (a) Lec2 and HEK293T-∆ST cells were incubated in HBSS buffer (pH 7.4) containing 3 mM HEPES, 20 mM MgSO_4_, the depicted concentration of CMP-Neu5Ac, and 40 μg/mL ST6Gal1 for 30 min at 37 °C. After three times washing with 1 × DPBS (pH 7.4, precolded at 4 °C), the cell-surface α2-6-linked sialic acids were detected with 20 μg/mL SNA-biotin. The resulted biotins were probed with 2 μg/mL Streptavidin-AF647 and quantified via flow cytometry. (b) The chemoenzymatically incorporated Neu5Ac-biotin on Lec2 cells or HEK293T-∆ST cells was probed by streptavidin-AF647 and quantified by flow cytometry. Error bars represent the standard deviation of three biological replicates.

**Figure S5.**
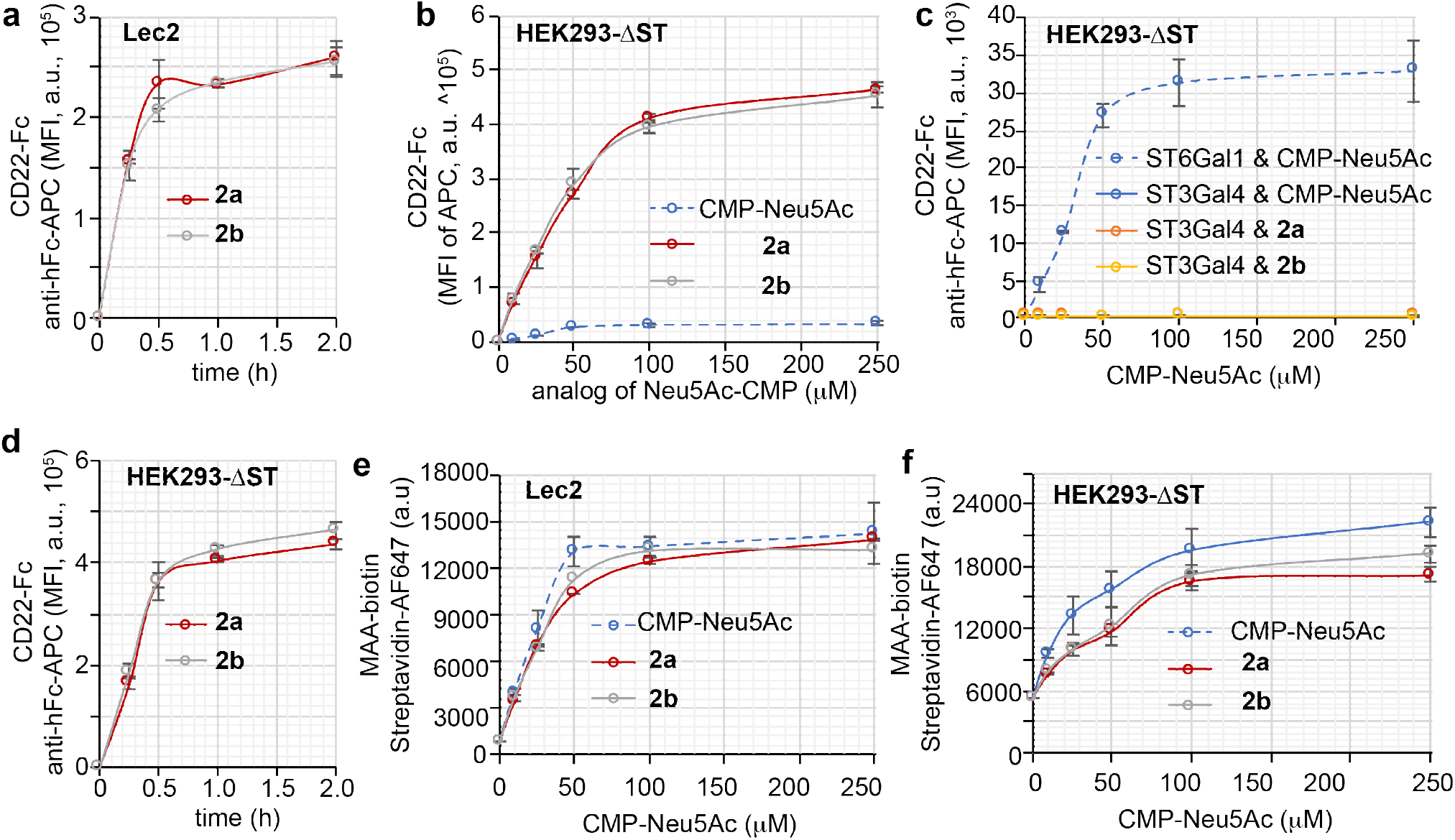
Chemoenzymatic incorporation of unnatural sialic acids with high affinity for human CD22. (a) ST6Gal1-assisted (40 μg/mL) incorporation of CMP-^BPC^Neu5Ac (**2a**, 100 μM) or CMP-^MPB^Neu5Ac (**2b**, 100 μM) onto Lec2 cells. (b and c) STs-assisted incorporation of the Neu5Ac analogs onto HEK293-∆ST cells. (d) Time-dependent incorporation of **2a** or **2b** onto HEK293-∆ST cells assisted by ST6Gal1. (e and f) ST3Gal4-assisted creation of α2-3-linked sialic acids on Lec2 cells (e) and HEK293-∆ST cells (f) via MAA-biotin staining. Error bars represent the standard deviation of three biological replicates.

**Figure S6.**
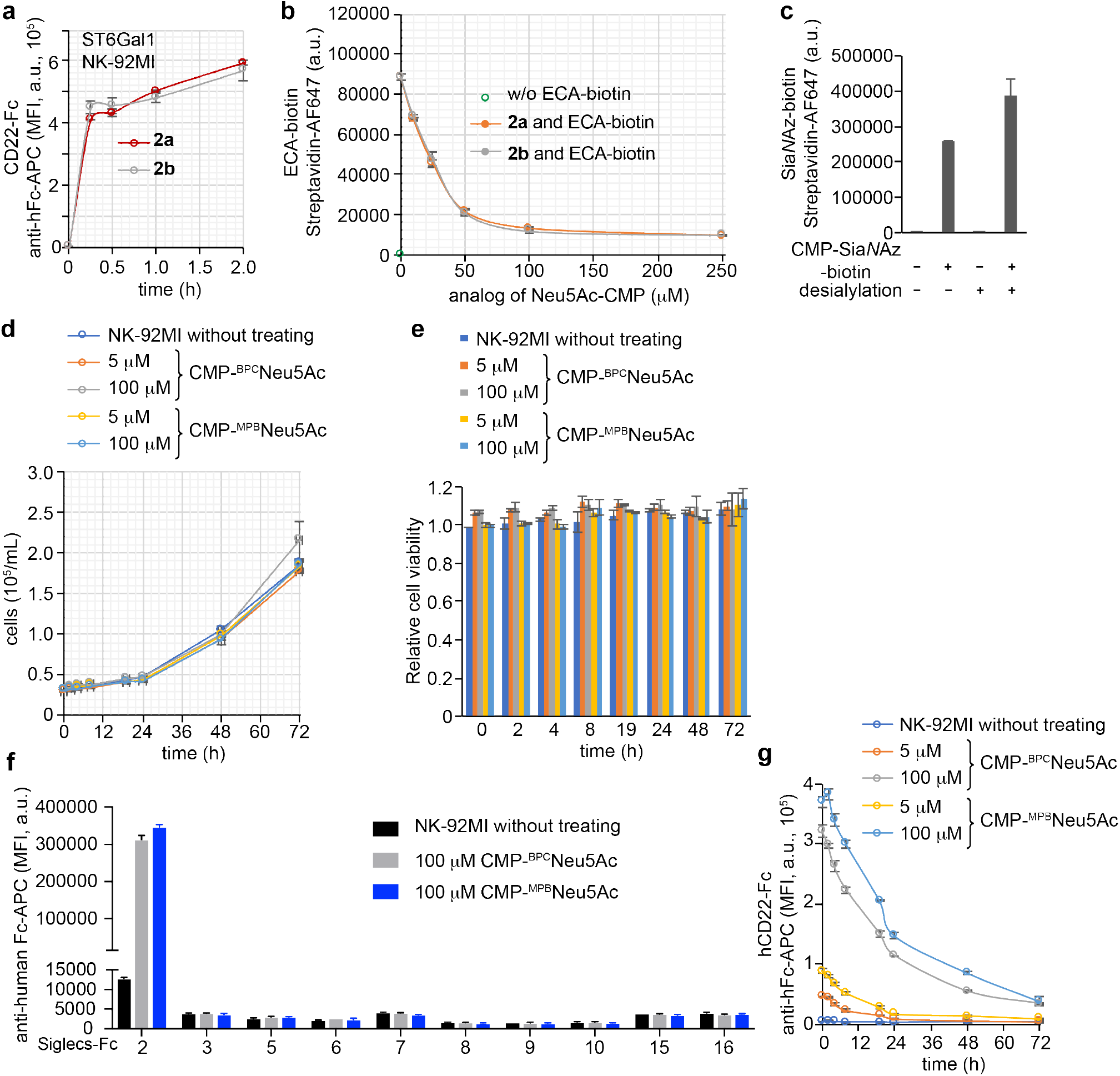
Chemoenzymatic incorporation of unnatural sialic acids with high affinity for human CD22 on NK-92MI cells. ST6Gal1-assisted (40 μg/mL) incorporation of CMP-^BPC^Neu5Ac (**2a**, 100 μM) or CMP-^MPB^Neu5Ac (**2b**, 100 μM) onto NK-92MI cells. (b) The incorporation of sialic acids on NK-92MI cell surface were confirmed via ECA-biotin staining (lectin specific for terminal Gal). (c) ST6Gal1-assisted (40 μg/mL) incorporation of CMP-Sia*N*Az-biotin (100 μM) onto NK-92MI cells pretreated with neuraminidase, or not. (c and d) In vitro proliferation (d) and cell viability (e) assessment of the NK-92MI cells with modifications or not. Cell proliferation was quantified by cell counts, and cell viability was assessed by DAPI staining. (f) NK-92MI cells with CD22 ligands or not were cultured in cell media and then stained with human CD22-Fc and anti-hFc APC antibodies before analyzing by flow cytometry at different time points post labeling. (g) The Siglec binding profiles of NK-92MI cells were assessed by Siglec-Fc binding. The *in situ* created glycoepitopes by ST6Gal1-assisted incorporation of CMP-^BPC^Neu5Ac (100 μM) or CMP-^MPB^Neu5Ac (100 μM) on NK-92MI cells showed significant specificity for human CD22 (Siglec-2). Error bars represent the standard deviation of three biological replicates.

**Figure S7.**
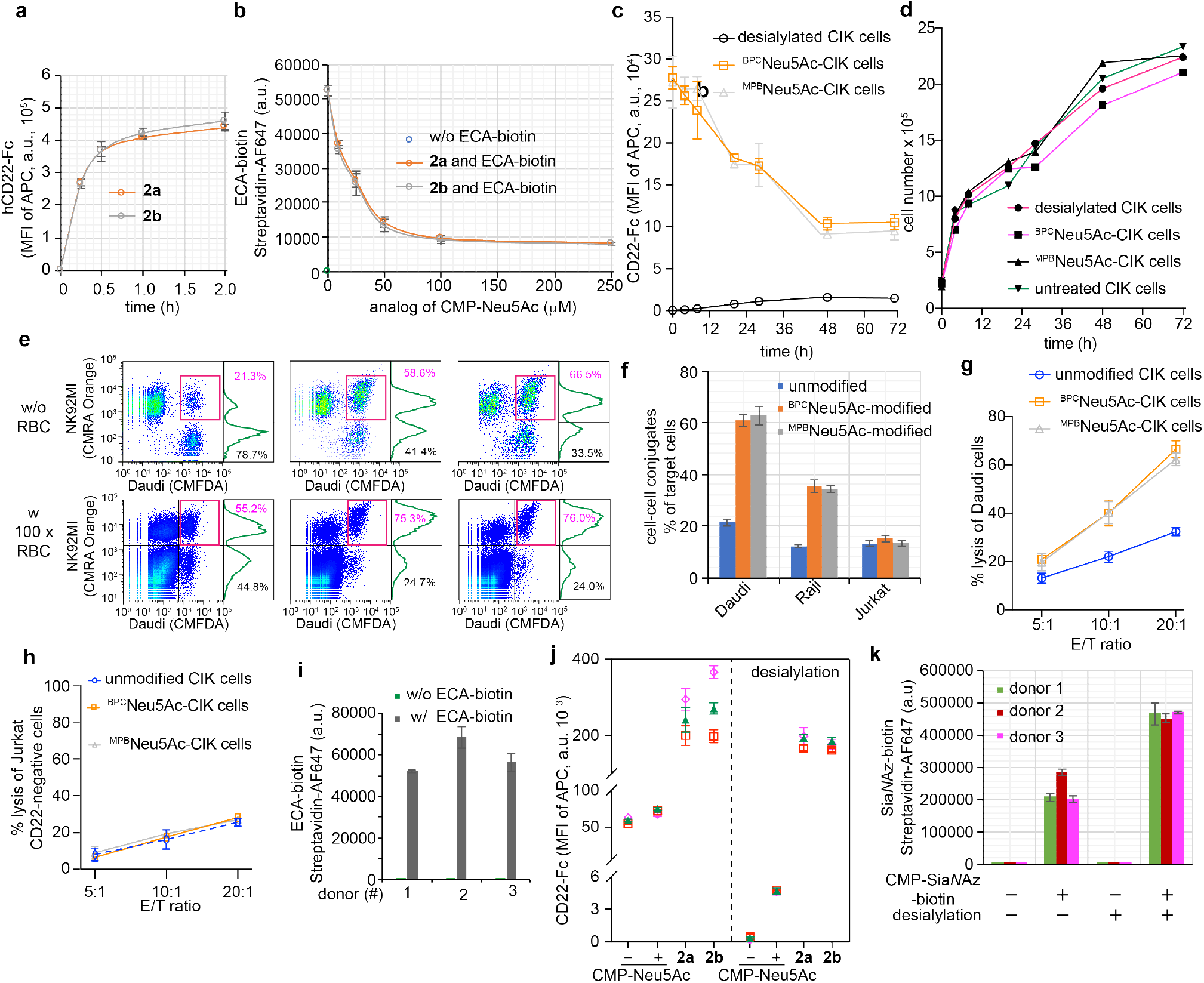
In situ creation CD22 high-affinity ligands on live cytokine induced killer (CIK) cells. (a) CD22-Fc profiling of the time-dependent (a) ST6Gal1-assisted incorporation of CMP-Neu5Ac and analogs on CIK cell surface. (b) The incorporation of sialic acids on NK-92MI cell surface were confirmed via ECA-biotin staining. (c) (d) Desialylation or incorporation unnatural sialic acids on CIK cells proliferated at a similar rate. (e) Flow cytometry-based detection of cell-cell conjugates of CIK cells and target cells (Daudi cells), in the presence of 100-fold RBC cells (first panel) or not (lower panel). (f) The relative population of CIK cell-cell conjugates of target cells. (g and h) LDH release assay quantifying cell-mediated cytotoxicity of CIK cells against Daudi (g), and Jurkat cells (g). (i) ECA-biotin based profile of the terimal Gal on the cell surface of CIK raised from three healthy donors (j) CD22-Fc profiling of the ST6Gal1-assisted incorporation of CMP-Neu5Ac and analogs on the cell surface of CIK raised from different healthy donors. (k) ST6Gal1-assisted incorporation of Sia*N*Az-biotin on CIK cells pretreated with neuraminidase or not before the glycocalyx engineering. In figures a-d, and f-k, the error bars represent the standard deviation of three biological replicates.

**Figure S8.**
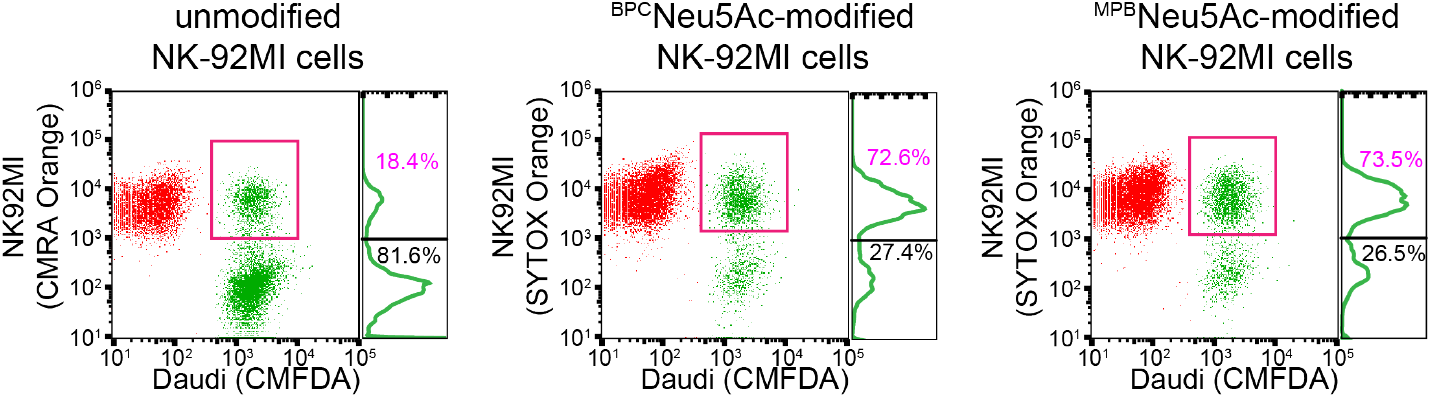
Ligand-dependent increase in B-lymphoma Daudi cell and NK-92MI cell-cell conjugates. NK-92MI cells modified or not were stained with cell tracker and incubated with Daudi cells at an E:T ratio of 5:1s. The NK-92MI and Daudi ‘cell-cell’ conjugates were analyzed by flow cytometry, and the numbers indicate the relative ratio of Daudi-lymphoma cells.

**Figure S9.**
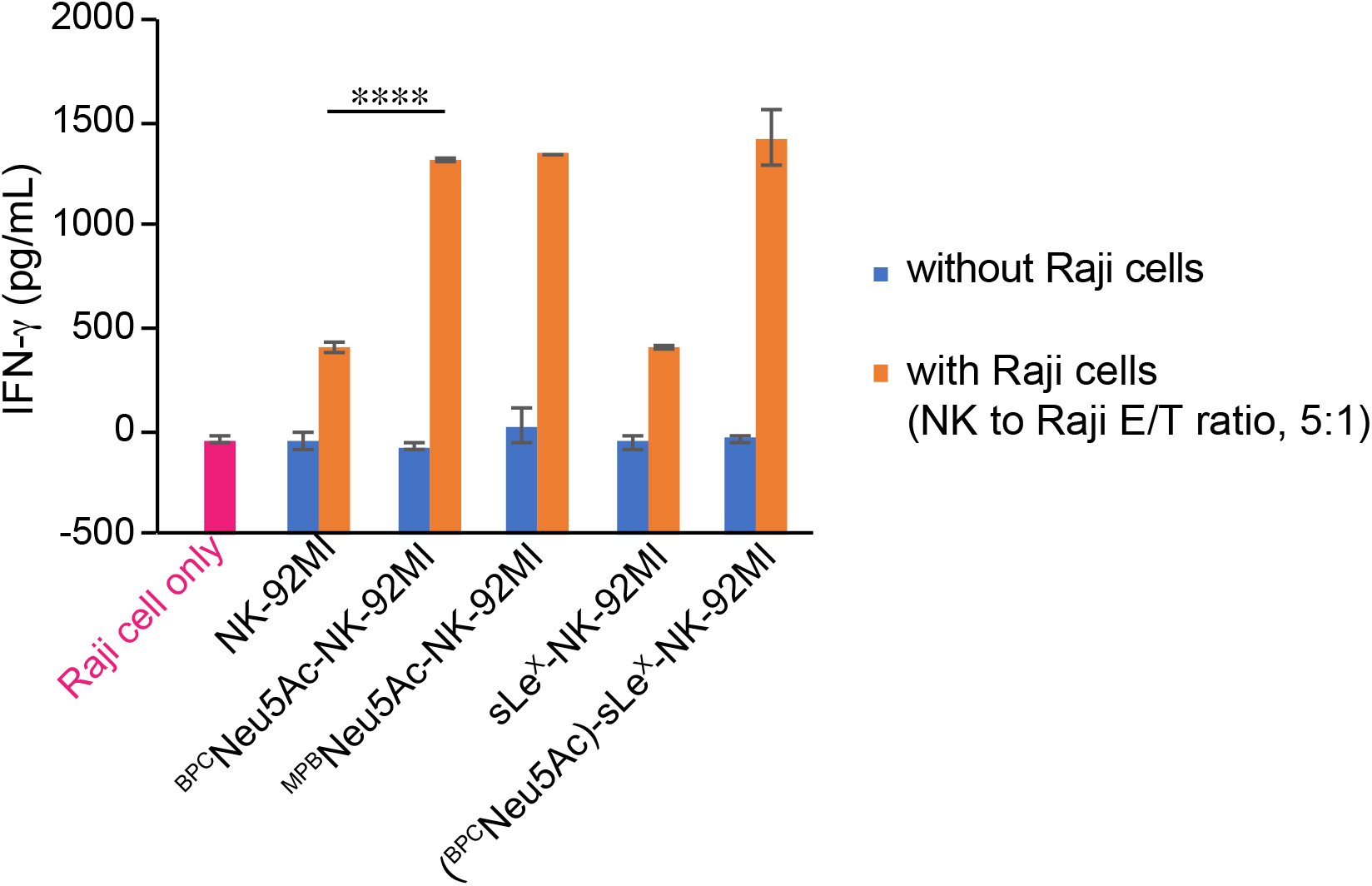
Ligand-dependent increase in NK-92MI cells’ IFN-gamma production. After glycosyltransferases-assisted engineering, NK-92MI cells were washed three times and incubated with Raji cells at an E/T ratio of 5:1 (about 5 × 10^5^ NK-92MI cells and 1 × 10^5^ Raji cells) for 1 hour. The supernatant of culture medium was collected and subjected to a human IFN-gamma quantification assay (Biolegend kit) following the supplier’s instructions.

**Figure S10.**
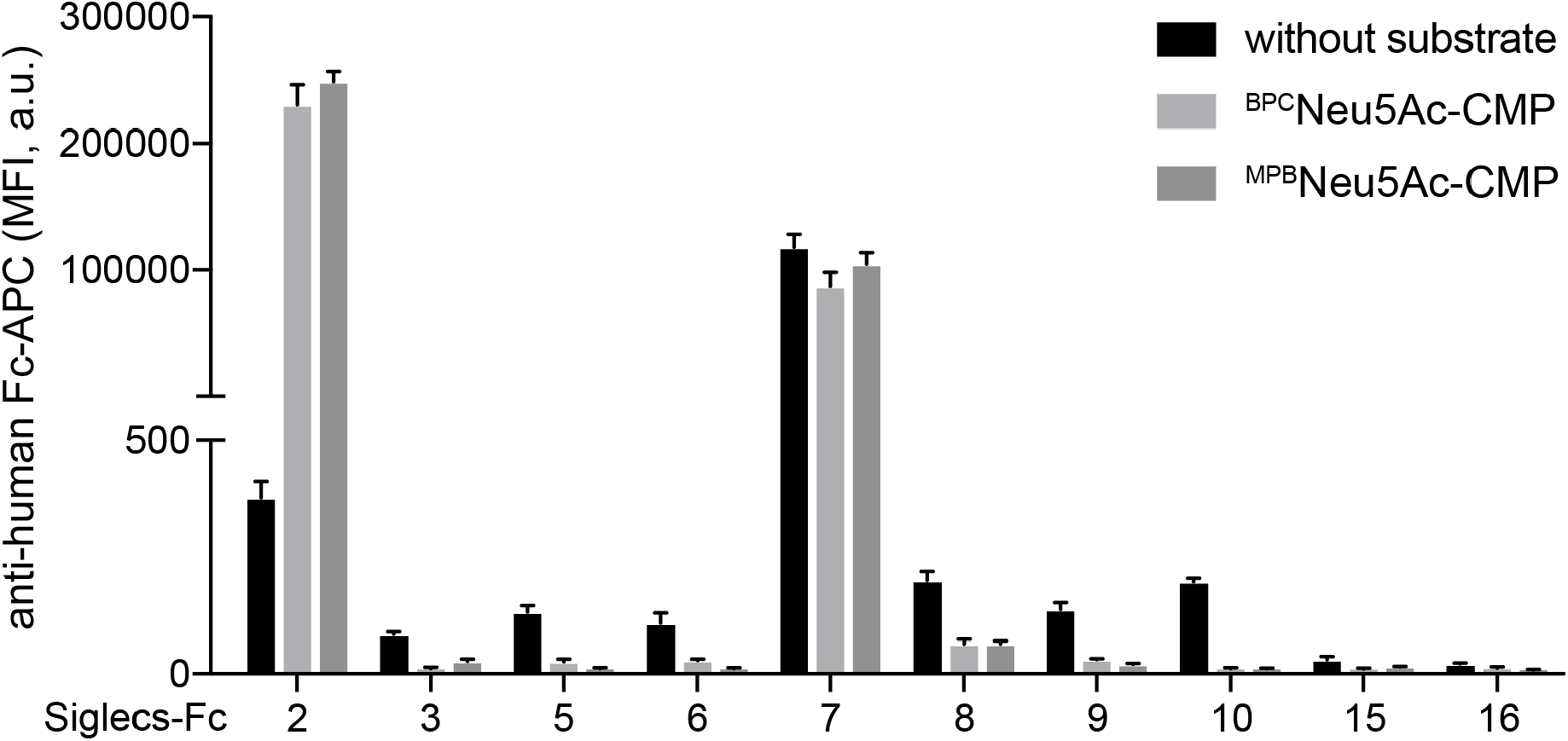
The incorporation of high-affinity human CD22 ligands on human red blood cell (RBC). The Siglec binding profiles of human RBC cells were assess by Siglec-Fc binding. The *in situ* creation of glycoepitopes by ST6Gal1-assisted incorporation of CMP-^BPC^Neu5Ac (100 μM) or CMP-^MPB^Neu5Ac (100 μM) on NK-92MI cells showed significant specificity for human CD22 (Siglec-2). Error bars represent the standard deviation of three biological replicates.

**Figure S11.**
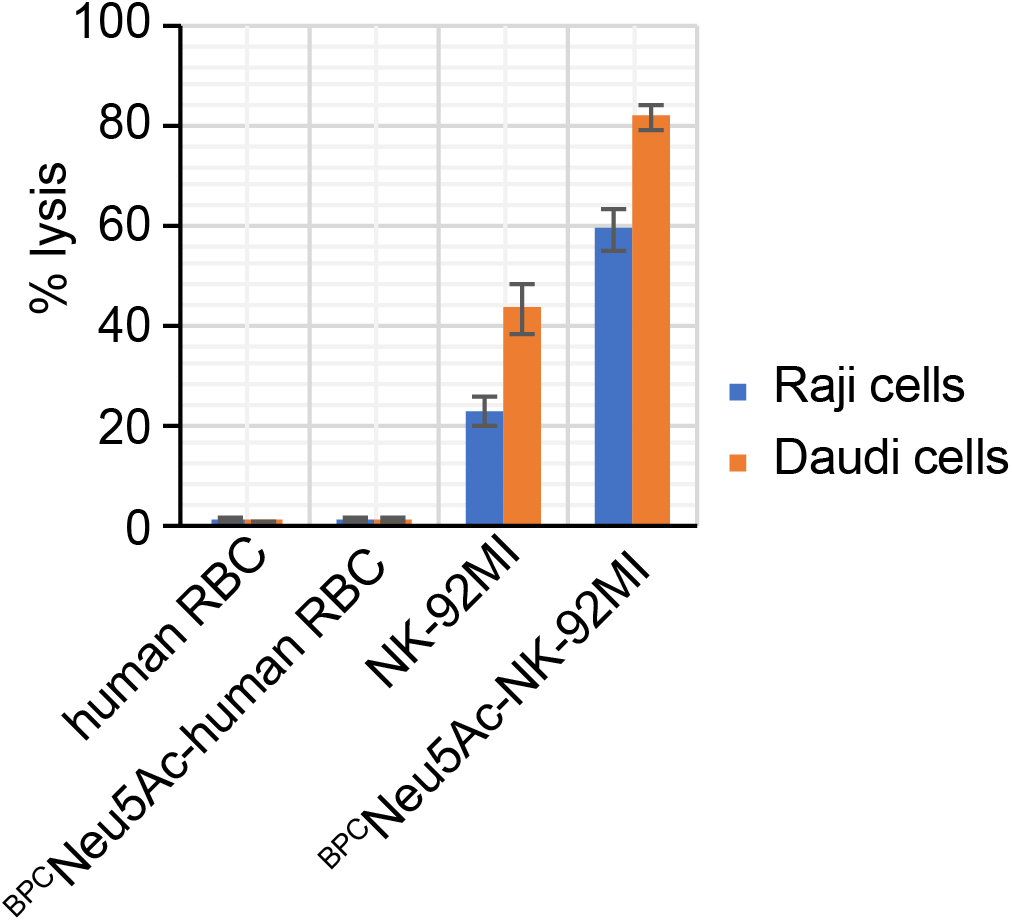
Assessing the cytotoxicity of the ligand-modified RBC towards B-lymphoma cells. Human RBC cells and NK-92MI cells modified with CMP-^BPC^Neu5Ac (100 μM) or not were incubated with B-lymphoma cells at an E:T ratio of 5:1 for 4 hours, including Daudi and Raji cells. The relative lysis of B-lymphoma cells was measured with an LDH kit. Error bars represent the standard deviation of three biological replicates.

**Figure S12.**
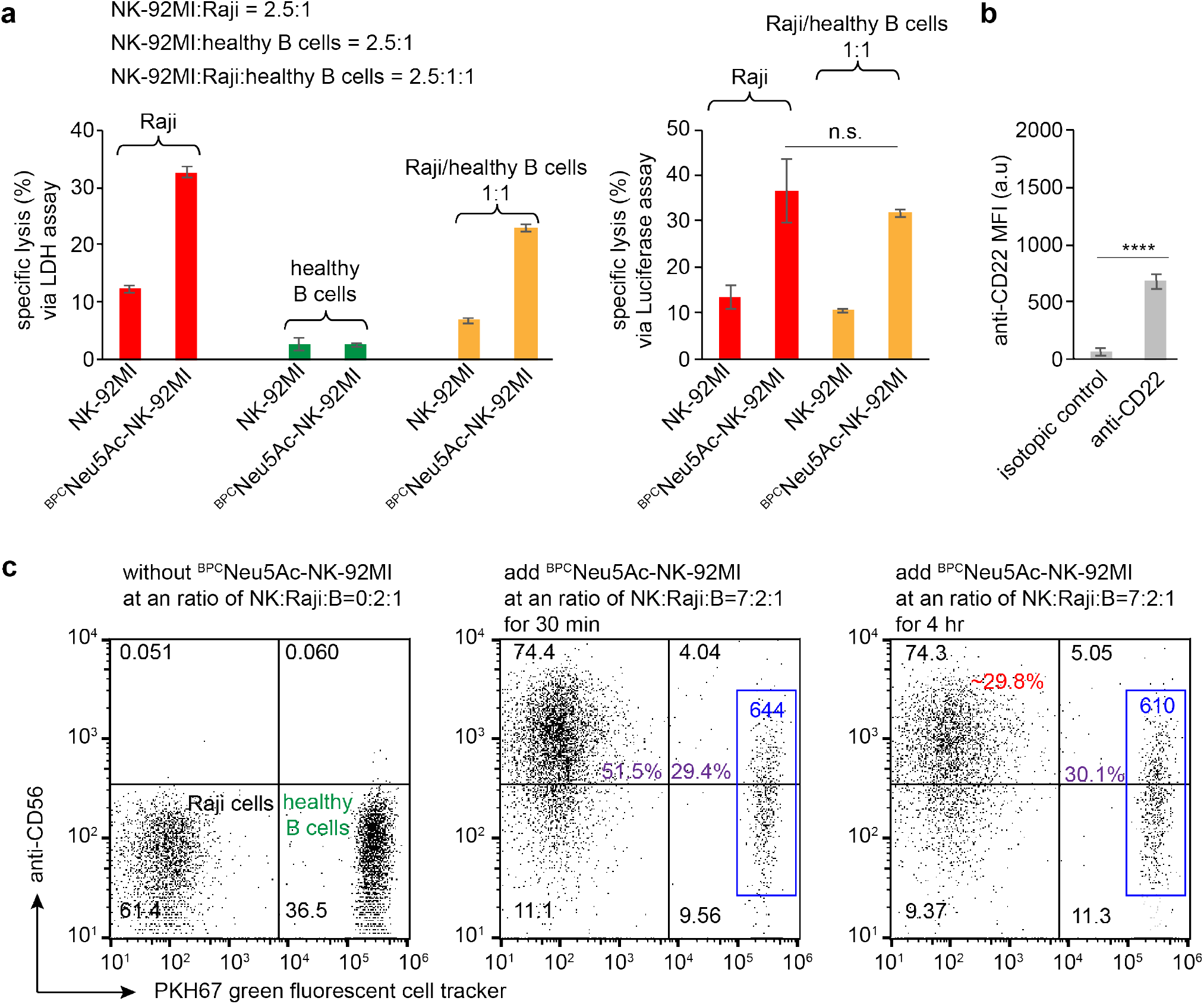
The incorporation of high-affinity human CD22 ligands on NK-92MI cells endowed CD22-specific killing of Raji cancer cells, but not the B-cells isolated from healthy human donor. (a) Specific lysis of NK-92MI cells with or without CD22-ligand loading (^BPC^Neu5Ac-NK-92MI) against target cells, including Raji cells, B-cells from healthy human donor. (b) CD22 antigen expression level of B-cells from healthy human donor. (c) Tracing ^BPC^Neu5Ac-NK-92MI cells selectively killing of Raji cells in coculture NK-92MI, Raji, and B-cells from healthy human donor. B-cells from healthy human donor were labeled with PKH67 cell tracker (Sigma), and NK cells were stained with anti-CD69 APC antibody. After 30-min incubation, we detected about 29.4 % and 51.5% of NK-target cell-cell conjugates of healthy B cells and Raji cells, respectively. Also, no dramatic decrease of healthy B-cell population was observed in 4-hr culture. The relative killing of Raji cells (red number show in three panel) was estimated based a similar ratio of NK-Raji conjugates and the ratio of Raji and healthy B cells in the begaining. In figure a and b, error bars represent the standard deviation of three experimental replicates.

**Figure S13.**
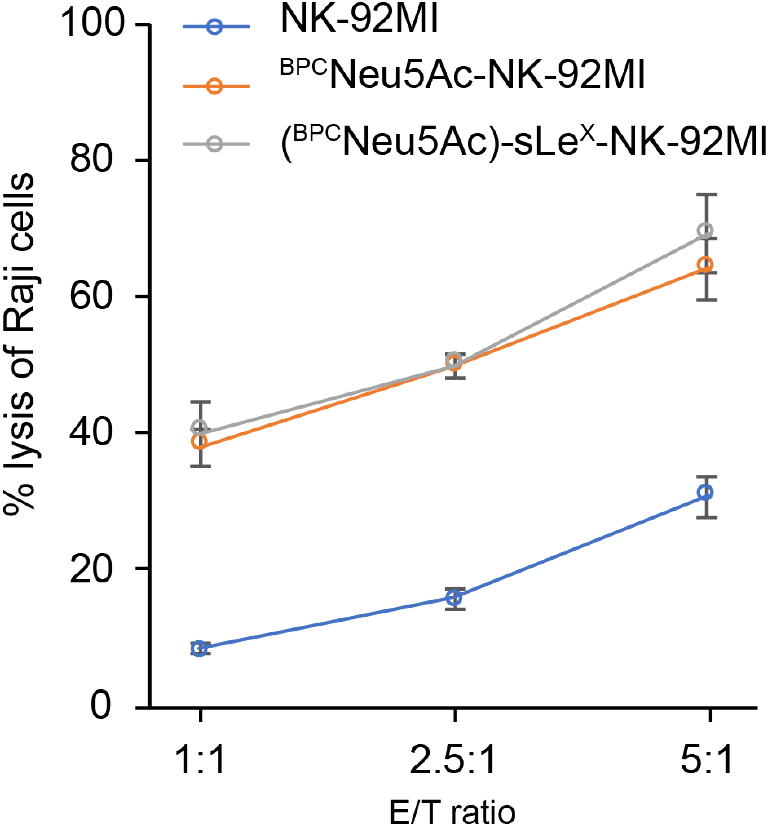
The cytotoxicity of siayl Lewis X engineered NK-92MI cells against Raji cells. NK-92MI cells were first treated with ST6Gal1 and CMP-^BPC^Neu5Ac (100 μM) for 30 min and washed three times with PBS. Secondly, the BPCNeu5Ac-loaded NK-92MI cells were further incubated with hFuT6 and GDP-Fuc (250 μM) for another 30 min to add fucose onto NK-92MI cells at α1-3-linkage. Error bars represent the standard deviation of three experimental replicates.

**Figure S14.**
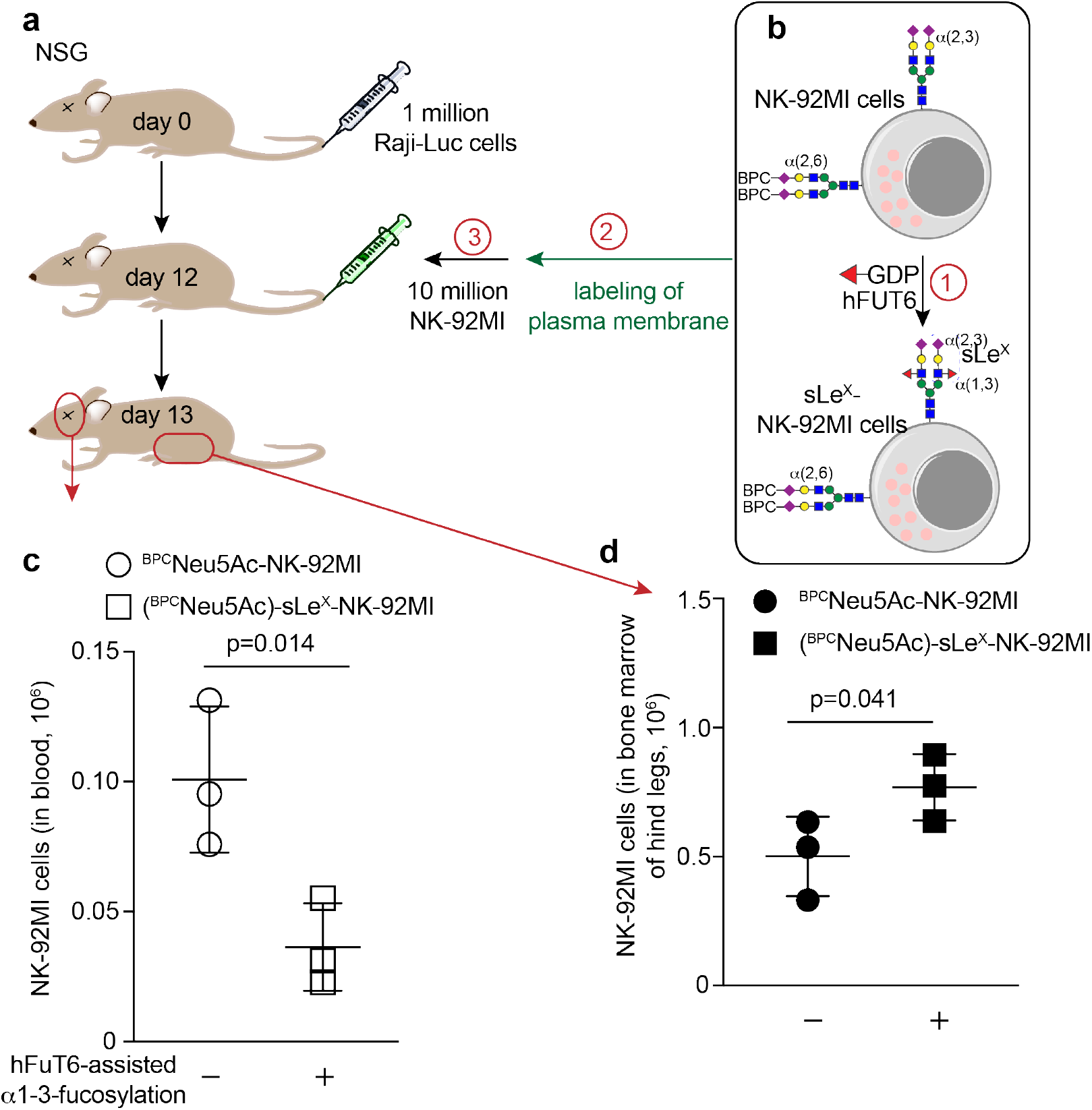
Plasma membrane tracker-assisted tracing of the adoptive NK-92MI cells in the Raji-Luc-inoculated NSG mice. On day 0, NSG mice were injected intravenously with 1 million Raji-Luc cells. 12 days later, the animals were then treated by i.v. injection of 10 million ^BPC^Neu5Ac-NK-92MI or (^BPC^Neu5Ac)-sLe^X^-NK-92MI cells. After overnight (about 10 hours) circulation, the blood and bone marrow were analyzed by flow cytometry. In figure c and d, p values were calculated via a two-sided Student’s t-test.

**Figure S15.**
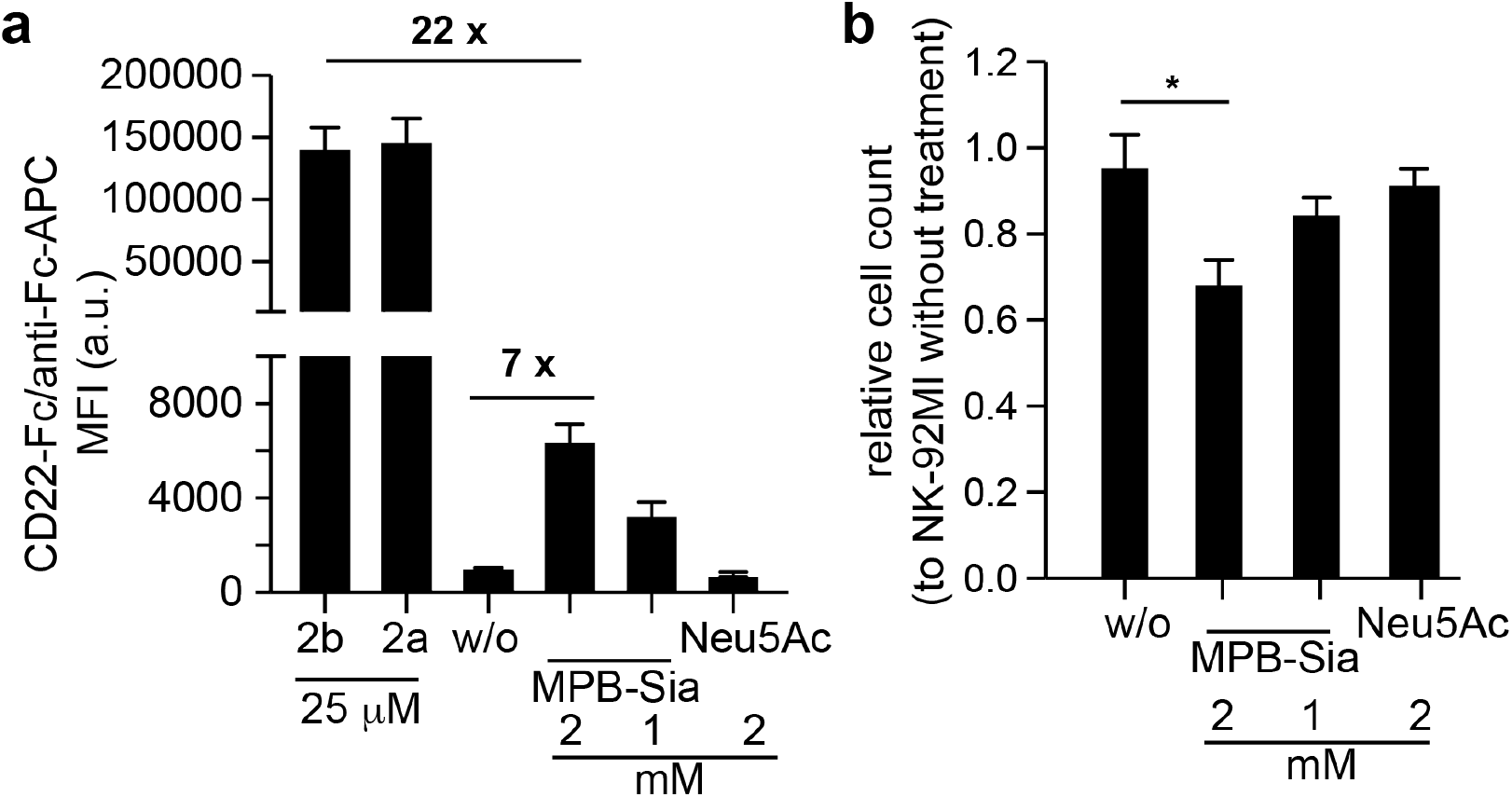
Comparision of high-affinity CD22 ligands presentation efficiency for CD22 targeting. To metabolic incorporation of ^MPB^Neu5Ac, the NK-92MI cells were cultured in media supported with 2 mM or 1 mM ^MPB^Neu5Ac, or 2 mM Neu5Ac, for 24 h. For ST6Gal1-assisted incorporation of ^MPB^Neu5Ac or ^BPC^Neu5Ac, the NK-92MI cells were treated with 40 μg/mL ST6Gal1 and 25 μM CMP substrates (CMP-^MPB^Neu5Ac (**2b**) or CMP-^BPC^Neu5Ac (**2a**)). (a) CD22-Fc binding profile of NK-92MI cells with or without cell-surface sialic acids engineering. (b) Cell growth of NK-92MI after 24 h sialic acids treatment. Error bars represent the standard deviation of three experimental replicates. In figure b, p value was calculated via a two-sided Student’s t-test and * indicate p<0.05.

**Figure S16.**
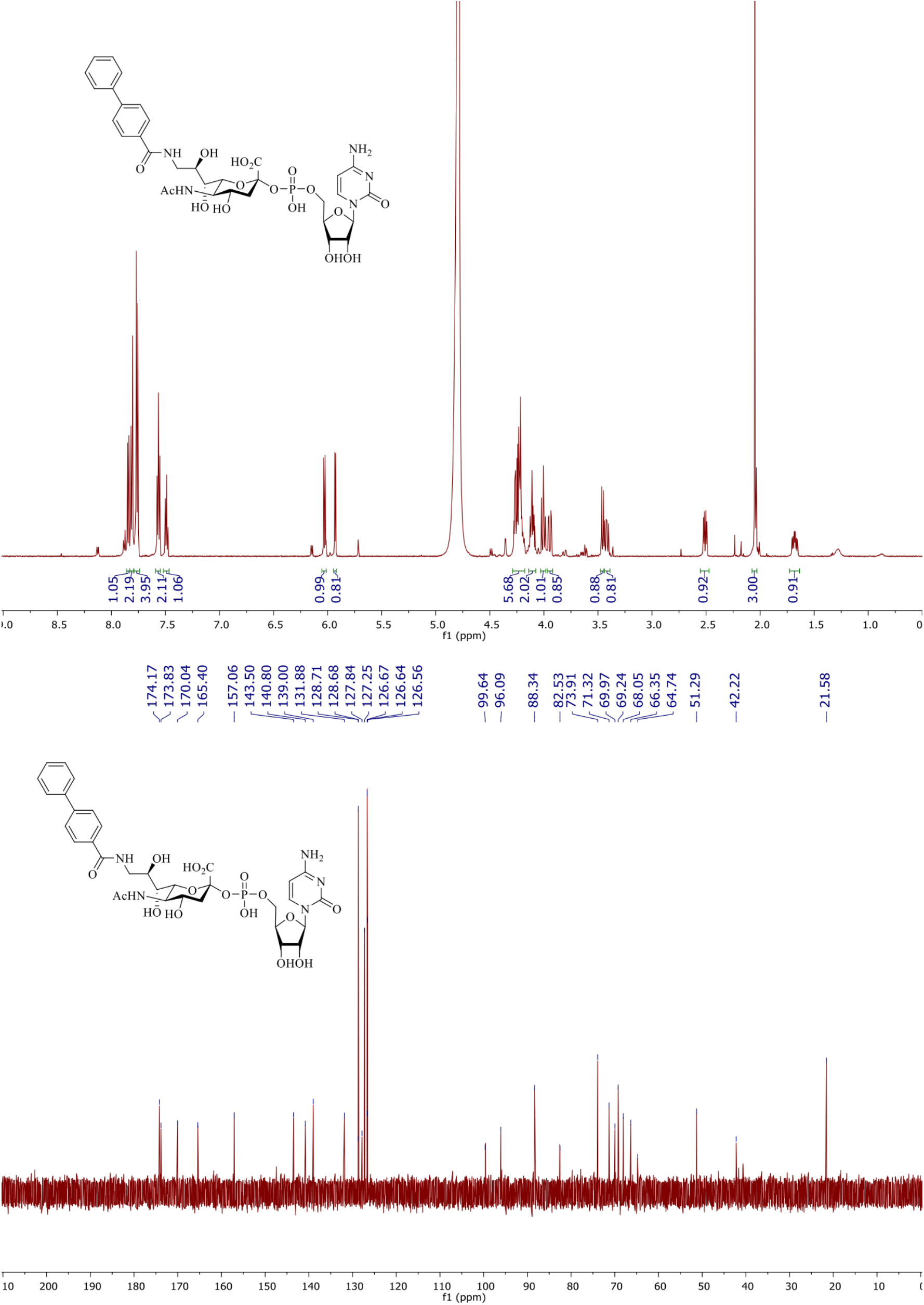
The NMR spectra of CMP-^BPC^Neu5Ac (****2a****). ^1^HNMR (600 MHz, D_2_O): *δ* = 1.68 (ddd, 1H, *J* = 6.0, 11.4, 13.2 Hz), 2.05 (s, 3H), 2.51 (dd, 1H, *J* = 4.8, 13.2 Hz), 3.42 (dd, 1H, *J* = 8.4, 13.8 Hz), 3.46 (d, 1H, *J* = 9.0 Hz), 3.95 (dd, 1H, *J* = 2.4, 13.8 Hz), 4.00 (t, 1H, *J* = 10.2 Hz), 4.08-4.14 (m, 2H), 4.18-4.29 (m, 6H), 5.93 (d, 1H, *J* = 4.8 Hz), 6.03 (d, 1H, *J* = 7.2 Hz), 7.47-7.51 (m, 1H), 7.54-7.58 (m, 2H), 7.74-7.86 (m, 7H). ^13^CNMR (150 MHz, D_2_O): *δ* = 21.58, 42.22, 51.29, 64.74, 66.35, 68.05, 69.24, 69.97, 71.32, 73.91, 82.53, 88.34, 96.09, 99.64, 126.56, 126.64, 126.67, 127.25, 127.84, 128.68, 128.71, 131.88, 139.0, 140.8, 143.5, 157.06, 165.4, 170.04, 173.83, 174.17. HRMS: m/z calc. for C_33_H_41_N_5_O_16_P: 794.2286; found: 794.2264 [M + H]^+^.

**Figure S17.**
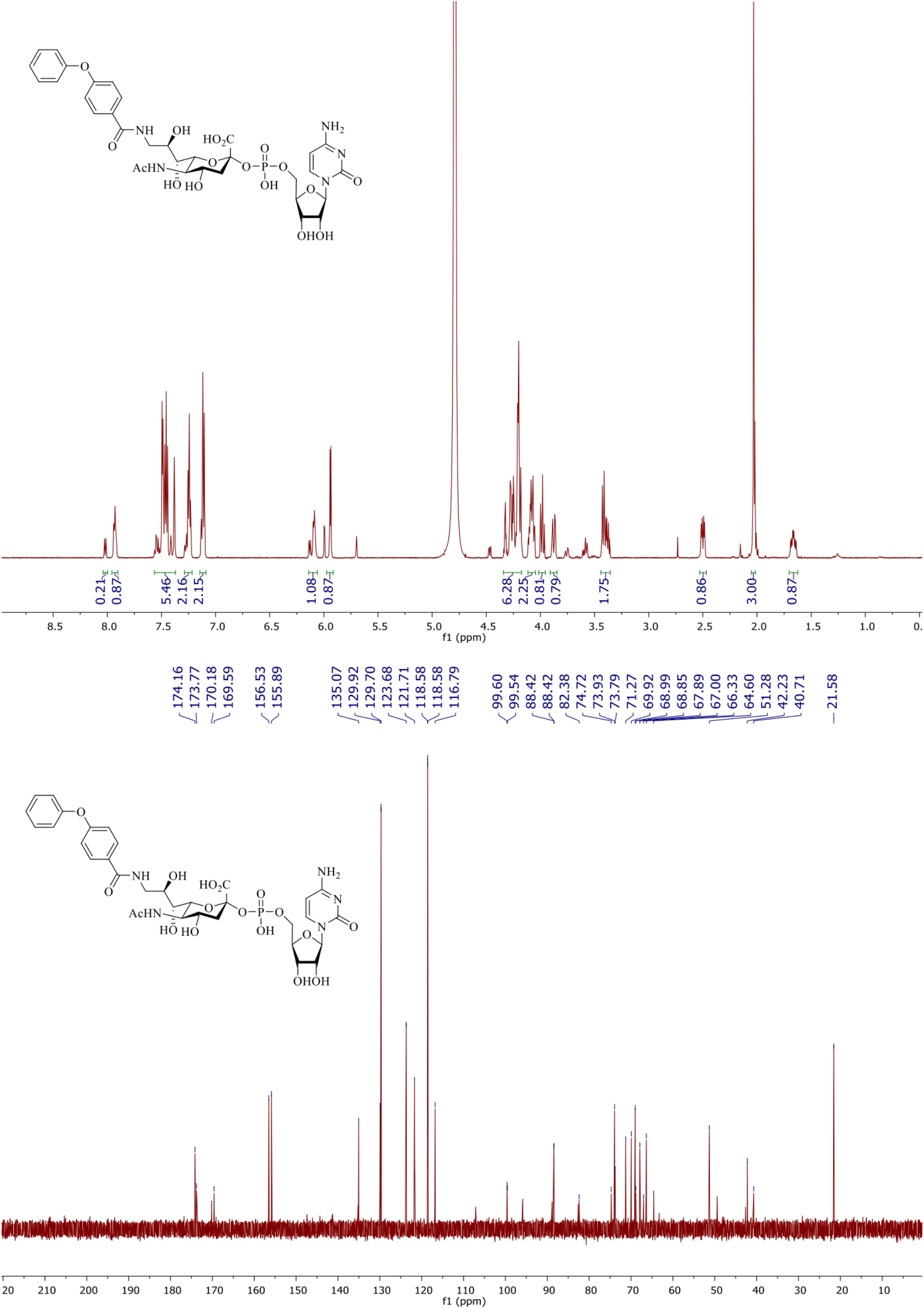
The NMR spectra of CMP-^MPB^Neu5Ac (**2b**) ^1^HNMR (600 MHz, D_2_O): *δ* = 1.66 (ddd, 1H, *J* = 6.0, 12.0, 13.2 Hz), 2.03 (s, 3H), 2.50 (dd, 1H, *J* = 4.2, 13.2 Hz), 3.35-3.44 (m, 2H), 3.88 (dd, 1H, *J* = 2.4, 13.8 Hz), 3.98 (t, 1H, *J* = 10.2 Hz), 4.04-4.12 (m, 2H), 4.17-4.34 (m, 6H), 5.94 (d, 1H, *J* = 4.8 Hz), 6.06-6.13 (m, 1H), 7.08-7.15 (m, 2H), 7.21-7.29 (m, 2H), 7.36-7.56 (m, 5H), 7.90-8.03 (m, 1H). ^13^CNMR (150 MHz, D_2_O): *δ* = 21.58, 40.71, 42.23, 51.28, 64.6, 66.33, 67.0, 67.89, 68.85, 68.99, 69.92, 71.27, 73.79, 73.93, 74.72, 82.38, 88.42, 88.42, 99.54, 99.6, 116.79, 118.58, 118.58, 121.71, 123.68, 129.7, 129.92, 135.07, 155.89, 156.53, 169.59, 170.18, 173.77, 174.16. HRMS: m/z calc. for C_33_H_41_N_5_O_17_P: 810.2235; found: 810.2250 [M + H]^+^.

## Supplementary Information

### Experimental Procedures

#### General methods and materials

All commercialized chemical reagents utilized in this study were purchased from suppliers, and used without further purification unless noted. Alkynyl-PEG_4_-biotin, Azide-PEG_4_-biotin and Al-kynyl-PEG-Cy3 were bought from Click Chemistry Tools (Scottsdale, AZ, USA); Streptavidin-Alexa Fluor 647 (AF488, AF594 and AF647) were obtained from Invitrogen (San Diego, CA, USA); Lectins (SNA-biotin and MAA-biotin) were purchased from Vector and used as instructed. Alexa Fluor 647-conjugated anti-human/mouse CLA antibody was purchased from Biolegend (cat. 321310). Horseradish peroxidase-conjugated anti-biotin antibody (HRP-anti-biotin antibody, 200-032-211) was purchased from Jackson ImmunoResearch Laboratories (West Grove, PA). The flow cytometry quantification was performed on an Attune NxT flow cytometer (Life). Absorbance, fluorescence intensity and luminescence were monitored in a Multi-Mode Microplate Reader (SynergyTM H4, Bio-Tek). Bioluminescence imaging of live mice was acquired using an IVIS Spectrum system.

#### Synthesis of chemical compounds

CMP-Neu5Ac^1^, GDP-fucose^2^, and CMP-Sia*N*Az-biotin^1^ were synthesized as previously reported. The CSS enzyme expression plasmid was obtained from Prof. Xi Chen, the enzyme was expressed and purified from *E. coli*.

General Procedure for preparation of the unnatural CMP-Sia.The corresponding unnatural sialic acid (0.1 mM), along with cytidine triphosphate (0.12 mM, 1.2 eq) were dissolved in 100mM Tris, 20 mM MgCl_2_ (3 mL) and the pH was adjusted to 8.5. 10 U of CMP-Neu5Ac synthetase was then added to the mixture. The reaction was kept in 37 °C for 2-4 h until TLC (ethylacetate : methanol : H_2_O = 5 : 4 : 1 v/v) indicated the reaction was complete. The reaction was then quenched with 5 mL methanol for 20 min and centrifuged at 4000 rpm to remove insoluble precipitates. The supernatant was evaporated to remove methanol and the remaining aqueous solution was frozen and lyophilized. After lyophilization, the residue was re-dissolved in H_2_O (1 mL). The solution was then loaded onto a P-2 column (1.5 × 100 cm) eluting with 50 mM NH_4_HCO_3_. The fractions containing product were combined, lyophilized and re-dissolved in H_2_O (1 mL). The solution was then vortexed, filtered through a 0.22 *μ*m filter and purified with HPLC.

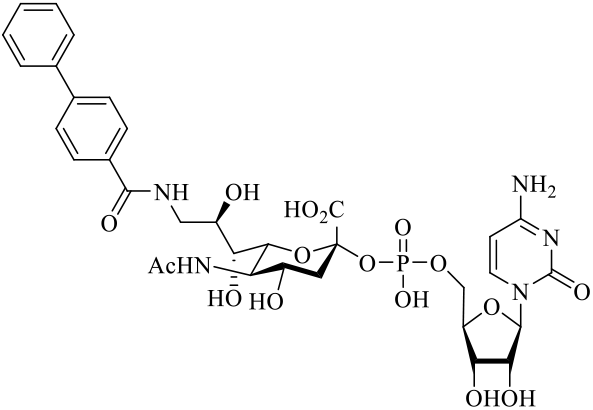

Compound **1a**^3^ was subjected to the general procedure for preparation of CMP-Sia, which gave compound **2a** at a yield of 78%.

^1^HNMR (600 MHz, D_2_O): *δ* = 1.68 (ddd, 1H, *J* = 6.0, 11.4, 13.2 Hz), 2.05 (s, 3H), 2.51 (dd, 1H, *J* = 4.8, 13.2 Hz), 3.42 (dd, 1H, *J* = 8.4, 13.8 Hz), 3.46 (d, 1H, *J* = 9.0 Hz), 3.95 (dd, 1H, *J* = 2.4, 13.8 Hz), 4.00 (t, 1H, *J* = 10.2 Hz), 4.08-4.14 (m, 2H), 4.18-4.29 (m, 6H), 5.93 (d, 1H, *J* = 4.8 Hz), 6.03 (d, 1H, *J* = 7.2 Hz), 7.47-7.51 (m, 1H), 7.54-7.58 (m, 2H), 7.74-7.86 (m, 7H). ^13^CNMR (150 MHz, D_2_O): *δ* = 21.58, 42.22, 51.29, 64.74, 66.35, 68.05, 69.24, 69.97, 71.32, 73.91, 82.53, 88.34, 96.09, 99.64, 126.56, 126.64, 126.67, 127.25, 127.84, 128.68, 128.71, 131.88, 139.0, 140.8, 143.5, 157.06, 165.4, 170.04, 173.83, 174.17. HRMS: m/z calc. for C_33_H_41_N_5_O_16_P: 794.2286; found: 794.2264 [M + H]^+^.

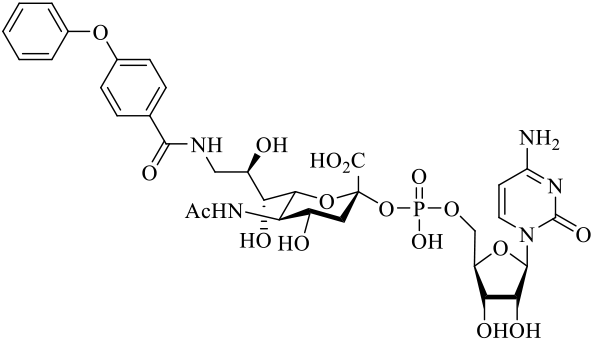

Compound **1b**^3^ was subjected to general procedure for preparation of CMP-Sia, which gave compound **2b** at a yield of 65%.

^1^HNMR (600 MHz, D_2_O): *δ* = 1.66 (ddd, 1H, *J* = 6.0, 12.0, 13.2 Hz), 2.03 (s, 3H), 2.50 (dd, 1H, *J* = 4.2, 13.2 Hz), 3.35-3.44 (m, 2H), 3.88 (dd, 1H, *J* = 2.4, 13.8 Hz), 3.98 (t, 1H, *J* = 10.2 Hz), 4.04-4.12 (m, 2H), 4.17-4.34 (m, 6H), 5.94 (d, 1H, *J* = 4.8 Hz), 6.06-6.13 (m, 1H), 7.08-7.15 (m, 2H), 7.21-7.29 (m, 2H), 7.36-7.56 (m, 5H), 7.90-8.03 (m, 1H). ^13^CNMR (150 MHz, D_2_O): *δ* = 21.58, 40.71, 42.23, 51.28, 64.6, 66.33, 67.0, 67.89, 68.85, 68.99, 69.92, 71.27, 73.79, 73.93, 74.72, 82.38, 88.42, 88.42, 99.54, 99.6, 116.79, 118.58, 118.58, 121.71, 123.68, 129.7, 129.92, 135.07, 155.89, 156.53, 169.59, 170.18, 173.77, 174.16. HRMS: m/z calc. for C_33_H_41_N_5_O_17_P: 810.2235; found: 810.2250 [M + H]^+^

#### Cell culture

Chinese hamster ovary (CHO) Lec 2, Daudi, Reh, Jurkat, and NK-92MI cells were obtained from American Type Culture Collection (ATCC, Manassas, VA). HEK293 cells with inactivated ST6Gal1, ST6Gal2, ST3Gal3, ST3Gal4, and ST3Gal6 (HEK293-∆ST) was a gift from Prof. Henrik Clausen (University of Copenhagen). Raji-Luc cells were a gift from Dr. Travis Scott Young (Calibr of Scripps), were obtained from ATTC by the Paulson group. Lec2 cells were routinely kept in half F12 medium and half high-glucose DMEM medium supported with 10% (vol/vol) heat-inactivated FBS, 100 U/mL penicillin G, and 100 mg/mL streptomycin. HEK293-∆ST cells were cultured in high-glucose DMEM medium supported with 10% (vol/vol) heat-inactivated FBS, 100 U/mL penicillin G, and 100 mg/mL streptomycin. NK-92MI cells were cultured in αMEM medium supported with 12.5% (vol/vol) heat-inactivated FBS, 12.5% (vol/vol) heat-inactivated horse serum, 100 U/mL penicillin G, and 100 mg/mL streptomycin. Cells were incubated in a water-saturated, 5% CO_2_ cultivator at 37 °C, free of mycoplasma contamination.

Primary human cytokine induced killer cell preparation: The human peripheral blood mononuclear cells (PBMC) were isolated from the blood of healthy donoars (200 mL blood samples per donor) via density gradient centrifugation; the PBMC cells were induced into CIK cells following former reports. All individuals provided informed consent for blood donation according to a protocol approved by the Internal Review Board and Ethics Committee. Briefly, diluted blood samples were layered into ficoll (Ficoll-Paque™ plus, GE healthcare)-preloaded SepMate^TM^-50 tubes (Stem cell). After centrifugation without breaking, the PBMC cells were collected, washed, and seeded in culture flasks at a concentration of 2 × 10^6^ cells/mL in RPMI-1640 medium (Gibco) containing 10% (v/v) FBS (Sigma), MEM NEAA (Gibco), and 10 mM HEPES (Gibco), 100 U/mL penicillin, and 100 U/mL streptomycin (Gibco). IFN-gamma (1000 U/mL) was added on day 0, the rhIL-2 (300 U/mL, Biolegend) and anti-CD3 (50 ng/mL, Biolegend) were added on day 1. CIK cell-expansion was performed for 3 weeks before being used for glycocalyx modfifications and targeting killing assays. Fresh medium and rhIL-2 (300 U/mL) were added every 2 days during culture, and the cell concentration was maintained at 2 × 10^6^ cells/mL.

#### Animal maintaince

All animal experiments were approved by the TSRI Animal Care and Use Committee. All the NSG male mice were bred or housed under specific pathogen free (SPF) conditions. NSG of breeders were purchased from Taconic Biosciences. NSG male mice 8–20 weeks of age were used for most experiments.

#### Chemoenzymaticy glycocalyx engineering via STs and CMP-Sia analogs

The truncated human recombinant STs, ST6Gal1 and ST3Gal4 were purified as previously reported.^4^ The cultured cells or primary cells were collected, washed twice with PBS, and resuspended in labeling buffer (HBSS buffer with 3 mM HEPES and 20 mM MgSO_4_). About 0.5 × 10^6^ cells were used in a total reaction volume of 50 μL containing ~2 μg enzyme and different concentrations of nucleotide sugar donors, including natural CMP-NeuAc and CMP-NeuAc analogs with high affinity for human CD22.

For lectin staining: After incubating at 37 °C for the depicted time periods, the cells were washed twice and resuspended in 50 μL HBSS buffer containing 10 mM CaCl_2_, 10 mM MgCl_2_, 10 μg/mL biotin-conjugated lectins (SNA-biotin and ECA-biotin), and 1 μg/mL DAPI. After being kept on ice in dark conditions for 30 mins, the cells were washed three times and resuspended in 100 μL HBSS buffer containing 10 mM CaCl_2_ and 10 mM MgCl_2_.

For one-step glycan labeling of cells, CMP-Sia*N*Az-biotin was used. After incubating at 37 °C for 30 mins, the cells were washed twice with PBS and resuspended in 50 μL FACS buffer (PBS containing 0.5 mM EDTA and 2 % FBS) with 5 μg/mL Alexa Flour 647-streptavidin and 1 μg/mL DAPI. Then, the cells were kept on ice in dark conditions for 30 min, washed twice and resuspended in 100 μL FACS buffer. After staining with AF647-streptavidin conjugates, cell surface fluorescence was detected by flow cytometry.

For Siglec-Fc staining: After incubating at 37 °C for the depicted time periods, the cells were washed twice with PBS and resuspended in 100 μL 30-min-premixed PBS buffer containing 2 μg/mL Siglec-Fc, 5 μg/mL anti-human IgG Fc-APC monoclonal antibody conjugates, and and 2% FBS. After a 30-min incubation, the cells were washed twice with PBS and resuspended in 150 μL FACS buffer (PBS containing 0.5mM EDTA and 2% FBS). The resuspended cells were then analyzed by flow cytometry. One-step biotin labeling of cell surface Lac*N*Ac containing glycans with Pm2,3ST and Pd2,6ST was performed in different cell lines.

#### Engineering sLe^X^ on NK-92MI cells via hFuT6

Truncated human recombinant hFuT6 was purified as previously reported.^4^ NK-92MI cells or ^BPC^Neu5Ac-modified NK-92MI cells (1 × 10^7^ cell/mL) were incubated in HBSS buffer (pH 7.4) containing 250 μM GDP-fucose, 40 μg/mL hFuT6, 3 mM HEPES, and 20 mM Mg^2+^ for 30 min at 37 oC. After washing three times with PBS, the in situ creation of sLe^X^ and α1-3-fucosylation were comfirmed with anti-sLeX antibody and AAL-biotin lectin, respectively. The cytotoxicity of the modified NK-92MI cells was assassed by in vitro killing of Raji cells, as well as IFN-gamma production. Then, modified NK-92MI cells were prepared for in vivo adoption.

#### Assessment of the NK-92MI cytotoxicity against B-lymphoma cells and CD22-negative Jurkat cells

Labeled or unlabeled NK-92MI cells were co-cultured with different types of cancer cells at the indicated effector/target ratios for 4 hours in a 96-well plate. Specific cancer cell lysis was detected by LDH secretion in supernatant (CytoTox96, Promega). Setup of control groups and calculations of specific lysis were done according to the manufacter’s instructions. The supernatant of each group (1-hour incubation) was also collected and subjected to IFN-gamma ELISA kits for quantification.

#### NSG mice model for assessing NK-92MI based killing of CD22-positive B-lymphoma cells

Male NSG mice (8-12 weeks old) were inoculated with 1 × 10^6^ Raji-Luc cells through tail vein injection (day 0). On day 2, mice were randomly divided into four groups and treated with HBSS, NK-92MI, ^BPC^Neu5Ac-modified NK-92MI, or ^BPC^Neu5Ac/sLe^X^ dual-labeled NK-92MI cells through tail vein injection (10 × 10^6^ NK-92MI cells per mouse). On day 5 after tumor challenge, mice were injected with 200 μL D-luciferin (15 mg/mL) through i.p. injection. Aftere 10 minutes, the bioluminescence signal in the mice was analyzed by the PerkinElmer IVIS system. The total photons indicating the tumor were quantified by the IVIS software. For tracing B-lymphoma progression, the NSG mice further received two additional NK-92MI doses on days 6 and 10. The luciferin-assisted IVIS monitoring of tumor growth were performed on day 5, day 7, day 12, and day 15. For the established tumor model, NSG mice were injected intravenously with 1 × 10^6^ Raji-Luc cells on day 0. The animals were imaged to confirm Blymphoma formation and received one treatment of 10 × 10^6^ NK-92MI or sLe^X^-NK-92MI cells (i.v. injection) on day 12.

#### Plasma membrane tracker based tracking of NK-92MI cells in tumor-inoculated NSG mice

To assess whether the tissue residency of NK-92MI cells armed by the in situ creation of sLe^X^ glycoepitopes was enhanced, NK-92MI cells treated with or without hFuT6-assisted α1-3-fucosylation were labeled by a live-cell plasma membrance tracking kit (thermo) following the suppler’s instructions. The labeled NK-92MI cells were them adopted into NSG mice that had been pre-adopted with Raji-Luc for 12 days. After overnight treatment, the NSG mice were sacrificed and the labeled NK-92MI cells in the bone marrow or blood samples were directly analyzed by flow cytometry.

#### Data and software availability

The flow cytometry data were processed using FlowJo (v10.0.3). The statistical analysis was performed in Prime and Microsoft Excel. The raw data that supported the findings of this study are available from the authors by reasonable request.

